# The fate of bacterial secondary metabolites in the rhizosphere: *Streptomyces* degrades and feeds on cyclic lipopeptides produced by competitors

**DOI:** 10.1101/2023.07.27.550914

**Authors:** Augustin Rigolet, Anthony Argüelles Arias, Adrien Anckaert, Loïc Quinton, Sébastien Rigali, Deborah Tellatin, Pierre Burguet, Marc Ongena

## Abstract

Cyclic lipopeptides are key bioactive secondary metabolites produced by some plant beneficial rhizobacteria such as *Pseudomonas* and *Bacillus*. They exhibit antimicrobial properties, promote induced systemic resistance in plants and support key developmental traits including motility, biofilm formation and root colonization. However, our knowledge about the fate of lipopeptides once released in the environment and especially upon contact with neighboring rhizobacteria remains limited. Here, we investigated the enzymatic degradation of *Bacillus* and *Pseudomonas* cyclic lipopeptides by *Streptomyces venezuelae*. We observed that *Streptomyces* is able to degrade the three lipopeptides surfactin, iturin and fengycin upon confrontation with of *B. velezensis in vitro* and *in planta* according to specific mechanisms. *S. venezuelae* was also able to degrade the structurally diverse sessilin, tolaasin, orfamide, xantholisin and putisolvin-type lipopeptides produced by *Pseudomonas*, indicating that this trait is likely engage in the interaction with various competitors.Furthermore, the degradation of CLPs is associated with the release of free amino and fatty acids and was found to enhance *Streptomyces* growth, indicating a possible nutritional utilization. Thereby, this work stresses on how the enzymatic arsenal of *S. venezuelae* may contribute to its adaptation to BSMs-driven interactions with microbial competitors. The ability of *Streptomyces* to degrade exogenous lipopeptides and feed on them adds a new facet to the implications of the degradation of those compounds by *Streptomyces*, where linearization of surfactin was previously reported as a detoxification mechanism. Additionally, we hypothesize that lipopeptide-producing rhizobacteria and their biocontrol potential are impacted by the degradation of their lipopeptides as observed with the polarized motility of *B. velezensis*, avoiding the confrontation zone with *Streptomyces* and the loss of antifungal properties of degraded iturin. This work illustrates how CLPs, once released in the environment, may rapidly be remodeled or degraded by members of the bacterial community, with potential impacts on CLP-producing rhizobacteria and the biocontrol products derived from them.

## Main

Cyclic lipopeptides (CLPs) represent a prominent and structurally heterogeneous class of molecules among the broad spectrum of small bioactive secondary metabolites (BSMs) formed by some plant beneficial rhizobacteria such as *Pseudomonas* and *Bacillus*^1,2^. These amphiphilic compounds consist of a partly or fully cyclized oligopeptide linked to a single fatty acid. They have been shown to inhibit the growth of a large range of phytopathogens and elicit immune responses in the host plant, leading to an induced systemic resistance (ISR) against infection by microbial pathogens^3,4^. These traits are largely responsible for the biocontrol potential of some CLP-producing isolates used to reduce plant diseases in sustainable agriculture^3^. From an ecological perspective, antimicrobial CLPs also contribute to the weaponry developed by these plant-associated bacteria to harm or kill microbial competitors in the densely populated rhizosphere niche. Moreover, CLPs support key developmental traits such as motility, biofilm formation or root colonization^2,3,5^.

CLPs are quite efficiently produced both *in vitro* and under natural conditions and substantial amounts are presumably released in the surrounding environment^5–7^. These metabolites are considered as chemically stable compounds due to the closed structure of the peptide moiety, the alternation of D- and L-amino acids and due to the incorporation of non-proteinogenic residues^2^. These molecules may thus accumulate in the rhizosphere, impact microbial interactions and modulate the composition of soil microbiomes. However, some recent studies reported instability of CLPs in the soil or in synthetic communities^8–10^. Yet, the mechanisms underlying CLP degradation as well as the possible ecological outcomes resulting from the phenomenon are poorly described.

In this work, we wanted to investigate the possible degradation of CLPs by *Streptomyces* as soil competitor and more specifically by *S. venezuelae* known for its metabolic robustness, behavioral plasticity and extensive enzymatic arsenal^11^. We first confronted the natural isolate *Streptomyces venezuelae* ATCC 10712 (Sv) to *Bacillus velezensis* strain GA1 (Bv), an archetypical root-associated isolate that efficiently co-produces surfactin, iturin and fengycin as the three lipopeptide families typical of the *B. subtilis* group^12,13^. Bacteria were inoculated at distance on gelified root exudate-mimicking medium designed to reflect the nutritional context of the rhizosphere (Fig. 1a). Sv colonies were phenotypically similar in interaction compared to monoculture while Bv colonies displayed a polarized growth and altered motility close to Sv (Fig. 1a, Supp. fig. 1). UPLC-qTOF-MS metabolite profiling of the compounds extracted from the agar in the confrontation zone revealed a decrease in the abundance of the three *Bacillus* CLP families compared with monocultures (Fig. 1b and Supp. fig. 1b), along with the accumulation of their cognate linearized forms eluting earlier (lower apparent hydrophobicity, Fig. 1b) and which were identified based on mass increment of 18 Da and MS/MS structure elucidation (Fig. 1c, Supp. fig. 2-4). Interestingly, additional ion species corresponding to shorter CLP fragments of surfactin (loss of the fatty acid from the linear form), iturin (loss of asparagine in position 3) and fengycin (loss of the terminal isoleucine) were also detected in the confrontation zone but not in Bv monoculture (Fig. 1b, MS/MS spectra in Supp. fig. 2-4). We next confronted Sv and the GFP-tagged GA1 upon colonization of tomato roots in a set-up better mimicking rhizosphere conditions. When inoculated alone, Bv readily colonizes roots as biofilm-structured colonies (Fig. 1d) and efficiently forms the three lipopeptides in their native cyclic structure as revealed by UPLC-MS analysis of rhizosphere extracts (Fig. 1e). Upon co-inoculation with Sv who forms mycelial pellets along the roots, there is no spatial exclusion of Bv, which still colonizes roots and secretes lipopeptides in substantial amounts. However, as for plate confrontation, a high proportion of linear iturins and surfactins (but not fengycins) along with surfactin fragment were observed in rhizosphere extracts indicating that some degradation of Bv CLPs by Sv also occurs under more natural settings of root co-colonization (Fig. 1e).

**Figure 1.**
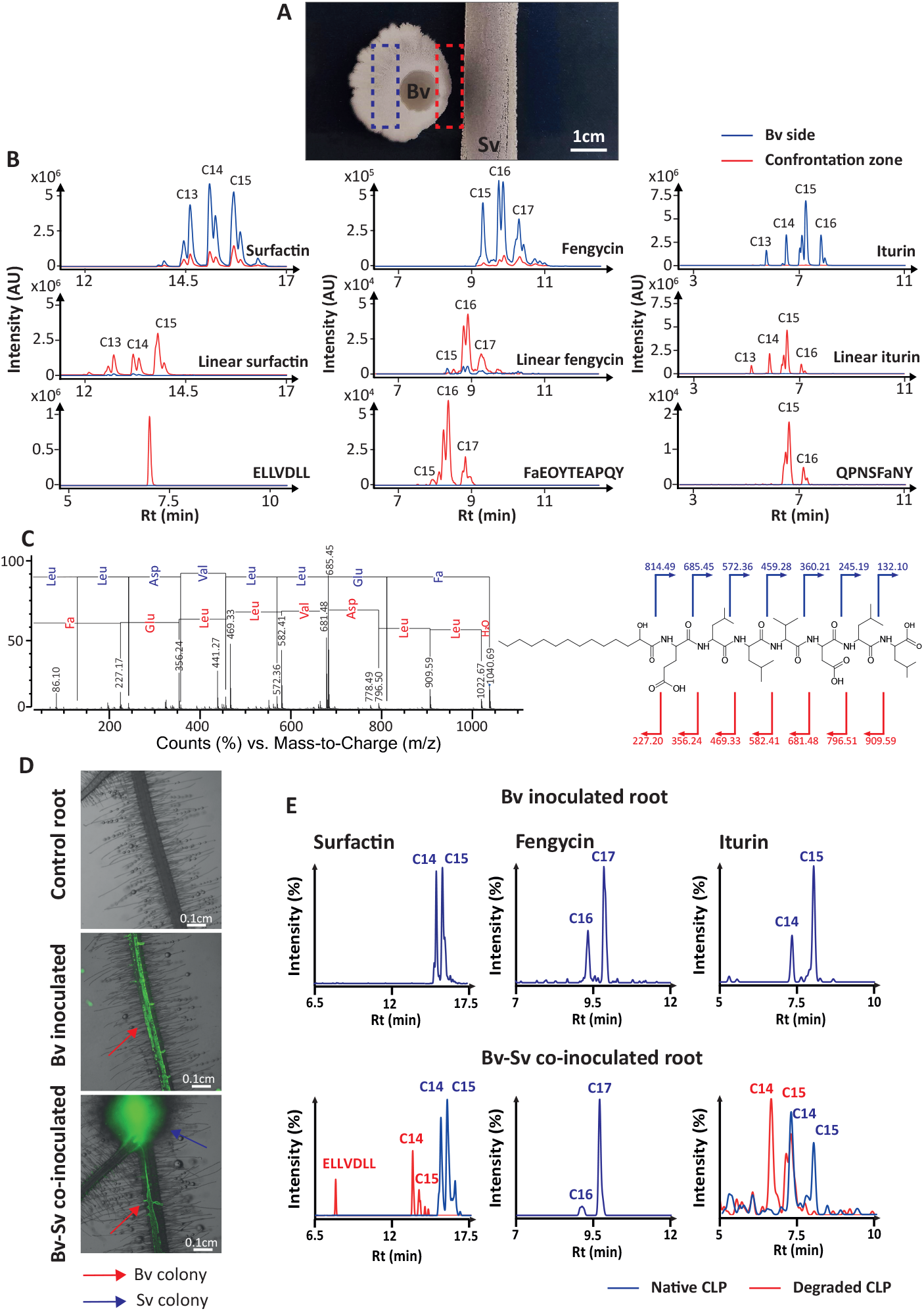
*Streptomyces venezuelae* ATCC 10712 linearizes *Bacillus* lipopeptides surfactin, iturin and fengycin upon interaction between *Bacillus velezensis* GA1. **a.** Picture of the interaction between *B. velezensis* GA1 (Bv, left side) and *S. venezuelae* ATCC 10712 (Sv, right side) on plate. Dashed rectangles represent the sampling areas used for metabolites extraction. The picture is representative of 3 biological replicates. **b**. UPLC-ESI-MS EIC of canonical, linear and degradation products of surfactin, iturin and fengycin extracted from agar in the interaction zone in-between Sv and Bv (confrontation zone, in red) and on the Bv side (in blue). The EIC are merged chromatograms of the [m+H]^+^ monoisotopic adducts of the main variants of each CLP. “Cn” represents the number of carbon of the fatty acid of the main CLP variant detected in each peak. Chromatograms are representatives of 3 biological replicates. Mean peak areas of the replicates of the different CLPs and linearized CLPs are shown in Supp. fig. 1. **c**. LC-ESI-MS/MS spectra of linear surfactin C14 and corresponding structure. Blue and red clippers and arrows represent the y- and b-ions. Fa stands for “fatty acid” **d**. Merged bright field and green fluorescens stereomicroscopic photos of tomato roots, not inoculated (control root, top picture), inoculated with GA1 GFPmut3-tagged (Bv inoculated, middle picture) and co-inoculated with GA1 GFPmut3 and *S. venezuelae* ATCC10712 (Bv-Sv co-inoculated, bottom picture). Pictures are representatives of 4 biological replicates. **e**. UPLC-ESI-MS EIC of canonical (blue) and linear (red) surfactin, fengycin and iturin extracted from tomato roots surrounding inoculated with Bv (top panels) and co-inoculated with Bv and Sv (bottom panel). Chromatograms are representatives of 4 biological replicates.

Based on these data, we further explored the Sv-mediated alteration of *Bacillus* CLPs and investigated the degradation process beyond linearization by using purified CLPs supplemented with Sv cell-free supernatant (CFS) on a time course experiment combined with feature-based molecular networking (FBMN). For each CLP, FBMN identified multiple degradation products generated in presence of Sv CFS, including those detected in confrontation assays and *in planta* (Fig. 2a,b,c, MS/MS spectra in Supp. fig. 2-4). Based on the fragments identified by FBMN and time-course monitoring of their occurrence (Fig. 2d, Supp. fig. 5), we propose a degradation mechanisms specific for each CLP characterized by the sequential generation of linearized lipopeptides followed by truncated fragments (Fig. 2a,b,c). In a similar set-up, we also tested Sv CFS for its ability to break down *Pseudomonas* CLPs representative of some of the main classes produced by soil-borne species^1^. Albeit to different degrees, sessilin, tolaasin, orfamide, xantholisin and putisolvin were all degraded (Supp. fig. 6-10) indicating that Sv may target a broad range of structurally diverse CLPs that the bacterium is likely to encounter in the soil. In most cases, degradation initiates with the opening of the peptide cycle followed by iterative degradation of the linear form, associated with the release of free fatty or amino acids. These mechanisms suggest the involvement of several enzymes secreted by Sv including esterase or endo-proteases for linearization and exo-proteases to further degrade the peptide. The enzymatic nature of the degradation was confirmed as heat-treated cell-free supernatant (CFS) of Sv completely loses its degradation activity (Supp. fig. 11) and comparative proteomic of active to inactive CFS of Sv highlighted the presence of several secreted proteases and amino acid/oligopeptide transporter unique to the active CFS of Sv (Supp. table 3).

**Figure 2.**
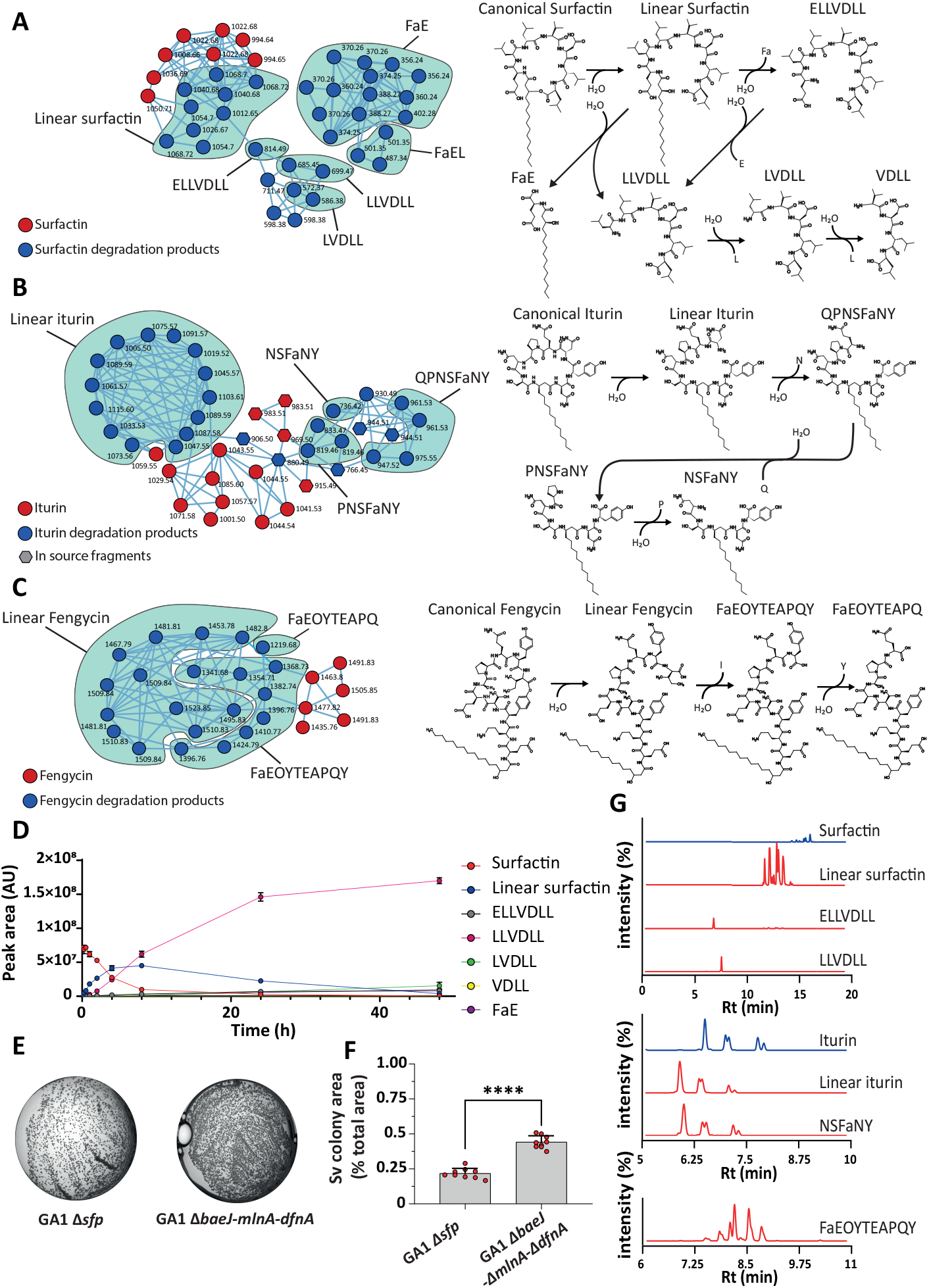
*S. venezuelae* degrades *B. velezensis* CLPs and feeds on it. **a, b, c**. Feature-based molecular networking of the degradation products of *Bacillus* CLPs surfactin (a), iturin (b) and fengycin (c) generated by Sv and proposed degradation mechanisms of each CLP. Pure CLPs were incubated for 24h 30°C at 100μM supplemented with 4% (v/v) of Sv CFS. MS/MS spectra are available in Supp. fig. 2-4. The degradation mechanisms are proposed based on the fragments detected. Summary of the identified features is available in Supp. table 4. **d**. Time course degradation of surfactin upon supplementation of Sv CFS 4% (v/v) **e**. Pictures of the Sv colony grown on gelified GA1 *ΔbaeJ-mlnA-dfnA* and GA1 *Δsfp* supernatant respectively. The *sfp* gene encodes for a 4’-phosphopantetheinyl transferase required for the activation of the synthesis of non-ribosomal peptides (NRPs) and polyketides (PKs). The mutant GA1 *Δsfp* is unable to synthesize the Sfp-dependent BSMs: the CLPs (surfactin, iturin and fengycin), the PKs (bacillaene, difficidin and macrolactin) and the siderophore bacillibactin. The mutant GA1 *ΔbaeJ-mlnA-dfnA* is unable to synthesize the PKs bacillaene, macrolactin and difficidin. Both strains were grown on iron sufficient medium to repress bacillibactin production. We used the mutants repressed in the synthesis of the PKs (*ΔbaeJ-mlnA-dfnA* and *Δsfp*) as Bv PKs inhibits Sv growth at high concentrations (i.e. when Sv grows on Bv CFS). Picture are representatives of 6 replicates. **f**. Relative summed colony area of Sv upon growth on GA1 *ΔbaeJ-mlnA-dfnA* and GA1 *Δsfp* supernatants respectively. Pictures areas used for colony area measurement =0.025cm^2^. Each dot represent a biological replicate (n=6). Statistical significance was calculated using Mann–Whitney test where (****: p< 0.0001). **g**. LC-ESI-MS EIC chromatograms of *Bacillus* CLPs iturin, surfactin and fengycin and the corresponding degradation products in Sv cultures grown on gelified GA1 *ΔbaeJ-mlnA-dfnA* CFS. The chromatograms are representatives of two biological replicates.

Hence, Sv can conceivably catabolize those exogenous CLPs and use them as nutritional sources. Indeed, we observed a significant increase in growth of Sv cultivated on gelified CLP-containing CFS of Bv mutants (GA1 *ΔbaeJ-mlnA-dfnA*, mutant unable to produce the three antibacterial polyketides bacillaene, difficidin and macrolactin, known for their toxicity toward *Streptomyces*^14^) compared to CLPs-free supernatants (GA1 *Δsfp*, mutant unable to produce the *sfp*-dependent metabolites: CLPs, PKs and the siderophore bacillibactin) (Fig. 2e,f). Extraction of the metabolites from the Sv cultures grown on CLP-containing conditions reveals the presence of degradation products of both iturin, surfactin and fengycin, further indicating that increased growth is driven by CLP catabolism (Fig. 2g). We propose that the ability of Sv to degrade CLPs and feed on them adds a new facet to the implications of CLPs degradation by *Streptomyces*, where linearization of surfactin was previously reported as a detoxification mechanism deployed by *Streptomyces* to counter the inhibition of aerial mycelium formation surfactin causes^15^. In environments marked by nutrient scarcity such as the rhizosphere, exogenous CLP degradation may thus represent a foraging strategy for Sv to access alternative sources of nutrients directly emanating from diverse microbial competitors.

Additionally, CLPs are key multifunctional BSMs whose biocontrol-associated activities often involve membrane perturbation and pore formation^16^. This CLP-membrane interaction is enabled by the peculiar amphiphilic 3D structures of those CLPs^17,18^. However, since the degradation alters their structures, it is likely associated to a loss of function. Indeed, *In vitro* experiments show that digested iturin loses its antifungal activities against phytopathogenic fungi *Fusarium* and *Botrytis in vitro* (Supp. fig. 12). Likewise, linear surfactin has been reported to lose its ISR triggering activity on tobacco cells^17,19^. Nonetheless, the impact degradation has on the biocontrol activities of other CLPs and on the biocontrol potential of CLPs-producing rhizobacteria deserves further investigation.

Furthermore, CLPs degradation also possibly hampers the producers as it may alter the promotion of phenotypical traits such as biofilm formation, motility and root colonization by those CLPs. The polarized motility of Bv away from Sv colonies observed in Fig. 1a may indeed result from the degradation of CLPs, especially surfactin, in the confrontation zone as it has been reported that structural modification of surfactin alters its ability to promote motility^20^. Yet, the actual impact of CLPs degradation on Bv phenotypes remains elusive.

Finally, the degradation of CLPs increases the chemical space resulting from the interaction as it generates numerous degradation products. Some of them may retain unsuspected bioactivities as recently reported. The degradation by *Paenibacillus* of the lipopeptide syringafactin produced by *Pseudomonas* generates toxic products to their common amoeba predators^21^ and the degradation of surfactin, also by *Paenibacillus*, serves as deterrent or territory marker in the interaction with *B. subtilits*^22^.

## Materials and Methods

### Strains and Cultures Conditions

#### Strains used are listed in Table S1

All experiments with *S. venezuelae* were inoculated with spores suspensions. *Streptomyces* spores were recovered from SFA medium plate (soy flour 20g/L, mannitol 20g/L, agar 20g/L, tap water 1L; pH 7.2) and stored at -80°C in peptone water (peptone 10g/L, NaCl 5g/L) supplemented with glycerol 25% (v/v). Spores concentration were measured with Bruker cells.

All *B. velezensis* GA1 wt and mutants and phytopathogenic bacteria were routinely precultured overnight in root exudates mimicking medium (REM; 0.5L *of all medium (0*.*685g of* KH_3_PO_4_, *21g of MOPS, 0*.*5g of* MgSO_4_.7H_2_0, 0.5g of KCl, 1.0g of yeast extract), 100μL of the trace solution (0.12g of Fe_2_(SO_4_)_3_, 0.04g of MnSO_4_, 0.16g of CuSO_4_ and 0.4g Na_2_MoO_4_ per 10mL) *and 0*.*5L of tobacco medium (*2.0g of glucose, 3.4g of fructose, 0.4g of maltose, 0.6g of ribose, 4.0g of citrate, 4.0g of oxalate, 3.0g of succinate, 1.0g of malate, 10g of fumarate, 1.0g of casamino acids, 2.0g of (NH_4_)_2_SO_4_ per liter, *Ph 7*.*0*) as described by ^23^ at 30 °C. After being washed three times in REM (cells were collected, centrifuged at 10 000rpm for 1 minute and resuspended in fresh REM), bacterial suspensions were set at proper OD_600nm_ and used for the experimental setup.

*Pseudomonas* strains were routinely precultured in casamino acid liquid medium (CAA; 10g/L casamino acid, 0.3g /L K_2_HPO_4_, 0.5g/L MgSO_4_ and pH 7.0), at 30°C. After being washed three times in casamino acid liquid medium (cells were collected, centrifuged at 10 000rpm for 1 minute and resuspended in fresh medium), bacterial suspensions were set at proper OD_600nm_ and used for the experimental setup.

### Construction of *Bacillus* Knock-out Mutant Strains

Triple mutant GA1 *ΔbaeJ-dfnA-mlnA* was constructed from GA1 *ΔbaeJ-dfnA* from Andric *et al*., 2022. On this mutant, *mlnA* gene was deleted by allelic replacement using a mutagenesis cassette containing a phleomycin resistance gene (50μg/mL) flanked by 1 kb of the upstream region and 1 kb of the downstream region of the targeted gene. Mutagenesis cassettes were constructed by overlap PCR as described by ^24^. The primers used were:

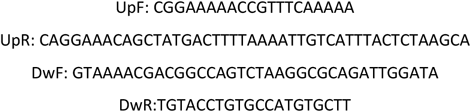

Recombination cassette was introduced in *B. velezensis* GA1 by inducing natural competence using a method adapted from ^25^. Briefly, after an initial preculture in LB medium at 37°C (160rpm) during at least 6h, cells were washed twice with peptone water. 1μg of the recombinant cassette was added to the GA1 cells suspension adjusted to an OD_600nm_ of 0.01 into MMG liquid medium (19g/L K_2_HPO_4_ anhydrous; 6g/L KH_2_PO_4_; 1g/L Na_3_ citrate anhydrous; 0.2g/L MgSO_4_ 7H_2_O; 2g/L Na_2_SO4; 50μM FeCl_3_ (filtrated on 0.22μm pore size filters), 2μM MnSO_4,_ 8g/L glucose, 2g/L L-glutamic acid, pH 7.0). After 24h of incubation at 37°C with shaking, double crossing over events were selected on LB plates supplemented with the adequate antibiotic. The gene deletion was confirmed by PCR analysis.

### Confrontation experiments

Confrontation assays were performed on square Petri dishes (12 × 12cm) with 40mL of REM solid medium at 26°C (REM supplemented with 14g/L of agar).

*S. venezuelae* ATCC10712 were inoculated as stripes (1× 12cm) in the middle of the plate with 40μL of spores suspension (10^7^ spores/mL) and spread with a cotton swab. Next, *B. velezensis* GA1 cells were collected from fresh precultures as described and adjusted to OD_600nm_ 0.1. Then, 5μL of *B. velezensis* GA1 suspension were spotted at 1cm of *Streptomyces* line. Control plates were done following the same procedure without the inoculation of either *Bacillus* or *Streptomyces*. Plates were then incubated for 3 days at 26°C in the dark. Pictures of the plates were then captured with a captured using a CoolPix camera (NIKKOR ×60 wide optical zoom extra-low dispersion vibration reduction [EDVR] 4.3 to 258 mm 1:33 to 6.5).

Metabolites accumulating in the vicinity of either *Bacillus* or *Streptomyces* and in the confrontation zone were recovered as followed: an area of agar (2 × 1cm) near the colony was sampled, transferred to Eppendorf tubes and placed for 24h at -20°C. Then, the agar were thawed at room temperature and centrifuged for 10 minutes at 13 000rpm. The supernatants were then collected and filtered (0.22μm pore size filters) before UPLC-MS analysis.

### In planta experiments

For the *in planta* studies, tomato seeds (*Solanum lycopersicum* var. Moneymaker) were sterilized following the protocol described by Hoff *et al*. (2021). Briefly, tomato seeds were primarily sterilized in 70% ethanol (v/v) by gently shaking for 2 minutes. Further, the ethanol was removed, and the seeds were added to the 50mL of sterilization solution (4.5mL of bleach containing 9.5% (v/v) of active chlorine, 0.01g of Tween 80, and 45.5mL of sterile water) and gently shaken for 10 minutes. Seeds were thereafter washed 10 times with water to eliminate sterilization solution residues. Sterilized seeds were then placed on square Petri dishes (12 × 12 cm) (5 seeds per plate) containing Hoagland solid medium (14g/L agar, 5mL of stock 1 [EDTA 5.20mg/L; FeSO_4_·7H_2_O 3.90mg/L; H_3_BO_3_ 1.40mg/L; MgSO_4_·7H_2_O 513mg/L; MnCl_2_·4H_2_O 0.90mg/L; ZnSO_4_·7H_2_O 0.10mg/L; CuSO_4_·5H_2_O 0.05mg/L; 1mL in 50mL stock 1, NaMoO_4_·2H_2_O 0.02mg/L 1mL in 50mL stock 1], 5mL of stock 2 [KH_2_PO_4_ 170mg/L], 5mL of stock 3 [KNO_3_ 316mg/L, Ca(NO_3_)_2_·4H_2_Omg/L], pH 6.5) and were placed in the dark to germinate for 3 days. Afterward, germinated seeds were inoculated with 2μL of the culture (OD_600nm_ 0.1) of *B. velezensis* GA1-GFP, 2μL of spore suspension (10^7^ spores/mL) of *S. venezuelae* ATCC10712 or with both GA1 and ATCC10712 (co-inoculation) and grown at 22°C under a 16/8h day/night cycle with constant light for 7 days.

For BSMs production analysis in *in planta* conditions, an agar part (1 × 1 cm) near the tomato roots was cut and weighted. Extraction and UPLC-MS analysis of the metabolites were then performed with the same protocol of the confrontation assays.

Stereomicroscopic pictures of inoculated tomato roots were taken with a Nikon SMZ1270 stereomicroscope (Nikon, Japan) equipped with a Nikon DS-Qi2 monochrome microscope camera and a DS-F 2.5× F-mount adapter 2.5x. Pictures were captured in the bright field channel and green widefield fluorescence (emission 535nm, excitation 470nm) with an ED Plan 2×/WF objective at an exposure time of 40ms. NIS-Element AR software (Nikon, Japan) was used to generate merged bright field and green fluorescence. Back ground and root green autofluorescence were removed by adjusting the LUTs (3388 to 6638)

### Generation of cell-free supernatants

The generation of cell-free supernatants (CFS) of Bv and Bv mutants (*Δsfp* or GA1 *ΔbaeJ*-*dfnA*-*mlnA*) were performed as followed. We inoculated 250mL flasks containing 50mL REM culture with at initial OD_600nm_ 0.02 with cells from fresh precultures as previously described. We then incubated the cultures for 24h at 30°C with continuous orbitally shaking (180rpm). Next, the cultures were centrifuged at 5000rpm at room temperature for 20 minutes. The supernatants were further filter-sterilized (0.22μm pore size filters) and stored at -20°C until further use..

Generation of *Pseudomonas* CFS was performed as followed: We inoculated 250mL flasks containing 50mL CAA culture with at initial OD_600nm_ 0.02 with cells from fresh precultures as described. We then incubated the culture for 48h at 26°C with continuous orbitally shaking (180rpm). Next, the cultures were centrifuged at 5000rpm at room temperature for 20 minutes. The supernatants were further filter-sterilized (0.22μm pore size filters) and stored at -20°C until further use.

*S. venezuelae* cultures were performed on ISP2 medium at 28°C (yeast extract 4g/L, malt extract 10g/L, glucose 4g/L, agar 20g/L; pH 7.3). On 12 × 12 cm Petri square plates, three stripes (1cm x 12) of spores were inoculated with a cotton swab (40μL of spores suspension (10^7^ spores/mL) each), spaced by 3cm each. The plates were left for 7 days of incubation. Next, the agar media was recovered and placed in Falcon tubes at -20°C for 24h. Then, the agar media were defrost at room temperature and centrifuged (8000rpm for 20 minutes). Finally the supernatant leaked from the agar was collected, filtered sterilized (0.22μm pore size filters) and stored at -20°C until further use. The heat treated Sv supernatant was generated by incubating an aliquot of the cell-free supernatant of Sv for 10 minutes at 98°C. The supernatant was then filtered (0.22μm pore size filters) and stored at - 20°C until further use.

### *In vitro* degradation of CLPs assays by *S. venezuelae*

The degradation assays of *Bacillus* CLPs by *S. venezuelae* were performed as followed: 500μL solutions of pure surfactin, iturin or fengycin (40μM) were supplemented by 4% (v/v) of *S. venezuelae* supernatant (or heat-treated supernatant) prepared as previously described. Then the solutions were incubated for 24h at 30°C with continuous shaking (180rpm). Next, the solution were centrifuged (1 minute at 10 000rpm), filtered (0.22μm pore size filters) and analyzed by UPLC-MS.

The degradation assays of *Pseudomonas* CLPs were performed using CFS generated as described above. They were supplemented by 4% (v/v) of *S. venezuelae* CFS prepared as described previously. Next, the solutions were incubated for 24h at 30°C with continuous shaking (180rpm). Then solution were centrifuged (1 minute at 10 000rpm), filtered (0.22μm pore size filters) and analyzed by UPLC-MS.

For the CLPs degradation kinetic experiments, surfactin, iturin and fengycin solutions supplemented with *S. venezuelae* supernatant were prepared following the same protocol. 20μL of solution were sampled at each time point and directly mixed with 80μL of acetonitrile to stop enzymatic degradation. Finally, they were stored at -20°C until analyzed by UPLC-MS.

### UPLC-MS analyses

All UPLC-MS analysis were performed using an Agilent 1290 Infinity II coupled with a diode array detector and a mass detector (Jet Stream ESI-Q-TOF 6530) in positive mode with the parameters set up as follows: capillary voltage of 3.5kV, nebulizer pressure of 35lb/in^2^, drying gas of 8L/min, drying gas temperature of 300°C, flow rate of sheath gas of 11L/min, sheath gas temperature of 350°C, fragmentor voltage of 175V, skimmer voltage of 65V, and octopole radiofrequency of 750V. Accurate mass spectra were recorded in the m/z range of 300 to 1,700. For untargeted MS/MS, we used the same MS1 parameters as described. We added MS2 untargeted acquisition mode with the parameters as follow: MS/MS range 50 to 1700m/z, MS/MS scan rate 3 spectra/s, Isolation width MS/MS medium (approx. 4amu), Decision Engine Native, Fixed Collision Energies 25V and 40V for surfactin, 50V for iturin and 60V for fengycin experiments, precursor selection : 3 for surfactin experiment, 4 for iturin and fengycin experiments, threshold 1500 (Abs), isotope model common, active exclusion after 2 spectra and released after 0.5 minute, sort precursors by charge state then abundance (charge state preference 1). For targeted MS/MS, we used the same MS1 parameters MS/MS range 50 to 1700m/z or 50 to 3200 (when required for *Pseudomonas* CLPs with mass >1700Da), MS/MS scan rate 3 spectra/s, Isolation width MS/MS narrow (approx. 1.3amu), Fixed Collision Energies 20, 40 and 60V. In all experiments, a C18 Acquity UPLC ethylene bridged hybrid (BEH) column (2.1mm × 50mm × 1.7μm; Waters, Milford, MA, USA) was used at a flow rate of 0.6mL/min and a temperature of 40°C. The injection volume was 20μL, and the diode array detector scanned a wavelength spectrum between 190 and 600nm. Otherwise mentioned, a gradient of acidified water (0.1% formic acid) (solvent A) and of acidified acetonitrile (0.1% formic acid) (solvent B) was used as mobile phase with a constant flow rate of 0.6mL/min, starting at 10% B and rising to 100% B in 20 minutes. Solvent B was kept at 100% for 4 minutes before going back to the initial ratio. MassHunter Workstation v10.0 and ChemStation software were used for data collection and analysis. For untargeted analysis of iturin and fengycin degradation products, we used the same solvent and flow rate, starting at 10% B to 20% in 2 minutes, then rising to 50% B at 14 minute and 100% B at 25 minute, followed by 6 minutes at 100% B and 5 at 10% B.

### MZmine-GNPS analysis

Mzmine 3 parameters used in this study are listed in supp. Table 3. Feature lists were then exported and submitted to GNPS. GNPS analysis of each CLP was performed with the parameter as follow: Quantification Table Source: MZmine, Precursor Ion Mass Tolerance: 0.02Da, Fragment Ion Mass Tolerance: 0.02Da, Min Pairs Cos: 0.5 for iturin and fengycin and 0.6 for surfactin, Minimum Matched Fragment ions: 4, Maximum shift between precursors: 500, Network TopK: 10, Maximum Connected Component Size: 100. All the other parameters were set as defaults. GNPS networks were then exported to cytoscape. Nodes corresponding to canonical CLP were identified based on the exact mass, the retention time, the presence in control CLP samples (without Sv supernatant treatment) and confirmed with the MS/MS spectra. Conversely, degradation products were identified as connected to canonical CLP and accumulating in CLP samples treated with Sv. The structures of the fragments were then elucidated with the MS/MS spectra.

The GNPS job ID are, for surfactin : ID=4c38af1675e744598573848474b784de, for iturin: ID=a2ac85a8fe49439fbf8235d514d31b1a and, for fengycin: ID=336c4c73ab6642f68787a173cf3ca719

### Growth on CLPs assays

The ability of *Streptomyces* to grow on *B. velezensis* CLPs were performed on 48 wells microplates. Each well was filled with 500μL of agar solution (40g/L agar) and 500μL of cell-free supernatant of *B. velezensis* GA1, GA1 *Δsfp* or GA1 *ΔbaeJ*-*dfnA*-*mlnA*. Wells were inoculated with 5μL of spore suspensions of *Streptomyces* (OD_600nm_ 0.1). The microplates were incubated for 3 days at 28°C. Stereomicroscopic pictures of inoculated tomato roots were taken with a Nikon SMZ1270 stereomicroscope (Nikon, Japan) equipped with a Nikon DS-Qi2 monochrome microscope camera and a DS-F 1× F-mount adapter 1x. Pictures were captured in the bright field channel and with an ED Plan 1×/WF objective at an exposure time of 20ms, gain 1.2X. NIS-Element AR software (Nikon, Japan) was used to generate bright field images. Colony area were measure by binary thresholding (LUTs <15100).

### Inhibition assays

For antifungal activities, we first prepared stock solution of spores. To that end, *Fusarium* and *Botrytis* fungi were grown on PDA (potato extract 4g/L, dextrose 20g/L, agar 15g/L) plates on the dark for 3 days at room temperature, followed by one day at daylight and 3 subsequent days in the dark. Spores were then collected, filtered and stored at -80°C in peptone water supplemented with glycerol 25% (v/v) for *Fusarium* and 50% (v/v) for *Botrytis*. The activity of iturin and linear iturin was quantified in microtiter plates (96-wells) filled with 250μL of LB liquid medium, inoculated with 10^6^ spores/mL of *Fusarium* or *Botrytis* from stock spores solutions at OD_600nm_ 0.01. The activities of iturin and linear iturin were estimated by measuring the pathogen OD_600nm_ every 30 minutes for 24h with a Tecan Spark microplate reader, continuously shaken at 150rpm and at 30°C.

## Acknowledgment

We warmly thank Guillaume Balleux and Pascale Bonnet for proof reading this document. We also gratefully acknowledge Sébastien Steels and Catherine Helmus for their technical support.

---

**Supp. fig. 1. Degradation of *Bacillus* lipopeptides surfactin, iturin and fengycin in interaction between *Bacillus velezensis* GA1 and *Streptomyces venezuelae* ATCC 10712. a**. Pictures of, from left to right, *Bacillus velezensis* GA1 alone (Bv), the interaction between *B. velezensis* (Bv, left side) and *S. venezuelae* (Sv, right side) and *S. venezuelae alone* (Sv). Dashed squares represent the sampling area for metabolites extraction. Pictures are representative of 3 biological replicates. **b**. Mean peak areas of canonical (blue) and linear (red) CLPs of *Bacillus* (surfactin, iturin and fengycin) extracted from agar in the control *Bacillus velezensis* GA1 (Bv), the control *S. venezuelae* (Sv) and in coculture of *B. velezensis* and *S. venezuelae*: on the side of *B. velezensis* (Bv side), in the interaction zone in-between Sv and Bv (confrontation zone) and on the *S. venezuelae* side (Sv side) as represented by the dashed squares in the pictures panel a. n=3 biological replicates. error bar indicating ± standard deviation. Peak area were measure from UPLC-ESI-MS EIC of the monoisotopic [m+H]^+^.

**Supp fig. 2. UPLC-ESI-qTOF MS/MS spectra surfactin degradation products generated by Sv**. Clippers represent B- and Y-ions sequences (in blue and red respectively). MS/MS spectra are merged spectra acquired at CID energy 20 and 40V.

**Supp. fig. 3. UPLC-ESI-qTOF MS/MS spectra of iturin degradation products generated by Sv**. Clippers represent B- and Y-ions sequences (in blue and red respectively). MS/MS CID energy was 50V.

**Supp. fig. 4. UPLC-ESI-qTOF MS/MS spectra fengycin degradation products generated by Sv**. Clippers represent B- and Y-ions sequences (in blue and red respectively). MS/MS CID energy was 60V.

**Supp. fig. 5. Degradation kinetics of surfactin, iturin and fengycin in presence of *S. venezuelae* supernatant**. Canonical CLP and degradation product content were measured by UPLC-ESI MS and are expressed as peak area. error bars represent the standard deviation n=3. Degradation kinetics were performed on 40μM pure surfactin, iturin and fengycin incubated at 30°C for 48h with 10% (v/v) filter-sterilized (0.22μM filters) *S. venezuelae* supernatant grown on ISP2 (adequate for enzyme production).

**Supp. fig. 6. Proposed degradation mechanisms of *Pseudomonas* spp. CLPs tolaasins, sessilins, orfamides, putisolvins and xantholysins by *S. venezuelae***. The mechanisms are inferred from the fragments detected and identified of each CLPs. To generate and identify degradation products, CFS of *Pseudomonas* containing the CLPs were incubated for 24 h at 30°C supplemented with 4% (v/v) of Sv CFS. The samples were the analyzed by UPLC-ESI-MS and structure were determined by UPLC-ESI-MS/MS. MS/MS spectra of the fragments are available in supp. Fig. 7-9.

**Supp. fig. 7. UPLC-ESI-qTOF MS/MS spectra orfamide degradation products generated by Sv**. Clippers represent B- and Y-ions sequences (in blue and red respectively). MS/MS CID energy was 20 and 40V. Red balls correspond to the positions of amino acid substitution in the orfamide variants

**Supp. fig. 8. UPLC-ESI-qTOF MS/MS spectra putisolvin degradation products generated by Sv**. Clippers represent B- and Y-ions sequences (in blue and red respectively). MS/MS CID energy was 20 and 40V. Red balls correspond to the positions of amino acid substitution in the putisolvin variants

**Supp. fig. 9. UPLC-ESI-qTOF MS/MS spectra Xantholysin degradation products generated by Sv**. Clippers represent B- and Y-ions sequences (in blue and red respectively). MS/MS CID energy was 20 and 40V. Red balls correspond to the positions of amino acid substitution in the Xantholysin variants

**Supp. fig. 10. UPLC-ESI-qTOF MS/MS spectra sessilin/tolaasin degradation products generated by Sv**. Clippers represent B- and Y-ions sequences (in blue and red respectively). MS/MS CID energy was 20 and 40V. Red balls correspond to the positions of amino acid substitution in the sessilin/tolaasin variants. Letters in blue in the peptide chain of the VSLVVQLVDhbTIHseDabK correspond to the amino acids in the peptide cycle

**Supp. fig. 11. The CLP-degradation activity of S. venezuelae is heat sensitive**. LC-MS EIC of canonical and linear iturins (left) and canonical and linear surfactin (right) in *B. velezensis* GA1 supernatant (top), *B. velezensis* GA1 supplemented with *S. venezuelae* ATCC10712 (middle) and *B. velezensis* GA1 supernatant supplemented with *S. venezuelae* ATCC10712 heat treated (10’ at 98°C) supernatant (bottom). Notation “Cn” correspond to the fatty acid chain length of the variant of surfactin and iturin associated to each peak. Y-axes of the chromatograms of the 3 conditions are linked for iturin and surfactin.

**Supp. fig. 12. Degraded iturin losses its inhibitory activity against fungal and bacterial phytopathogens**. Optical density of *B. cinerea* and *F. lactuca* liquid cultures supplemented with canonical or degraded iturin. Graphs show the mean optical density (OD) and ±SD calculated for 6 biological replicates (n=6). OD was measured on 96 wells microplates inoculated with 10^6^ spores/ml and grown for 96 and 48h for *Botrytis* and *Fusarium* respectively. Ctrl correspond to fungal culture without (linear) iturin supplementation. Statistical comparison between control and supplemented with CLPs was performed based on T-test (ns: not significant, *: p<0.05, **** p<0.0001).

**Supp. table 1. Stains used in this study**

**Supp. table 2**. MZmine parameters used for FBMN

**Supp. table 3. Secreted proteins found only in active Sv CFS with function possibly related to CLP catabolism**.

**Supp. table 4. Features identified in** Fig. **1 a**,**b**,**c**.

