## Supplementary Figures and Tables for "The fate of bacterial secondary metabolites in the rhizosphere: *Streptomyces* degrades and feeds on cyclic lipopeptides produced by competitors"

### Supp. Fig. 1

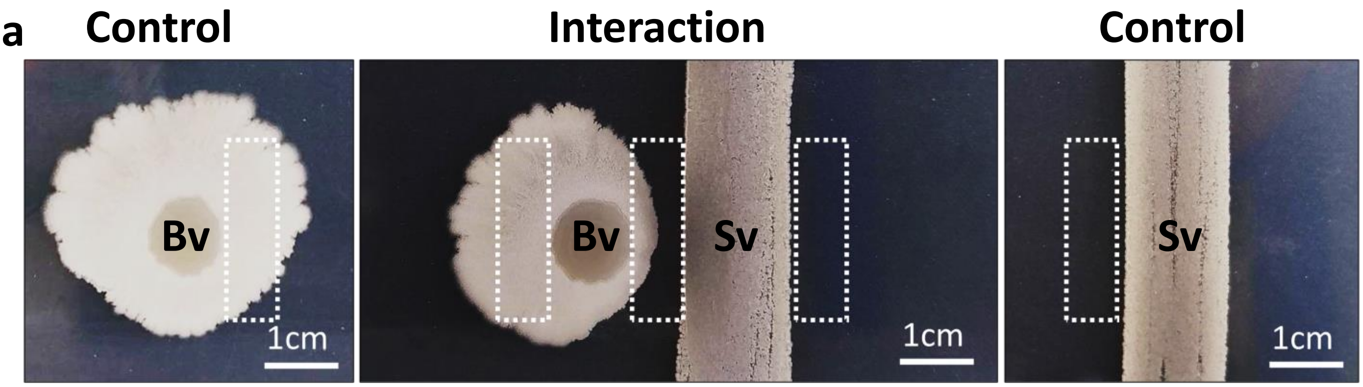

a
Control
Interaction
Control
Bv
Bv
Sv
Sv
1cm
1cm
1cm


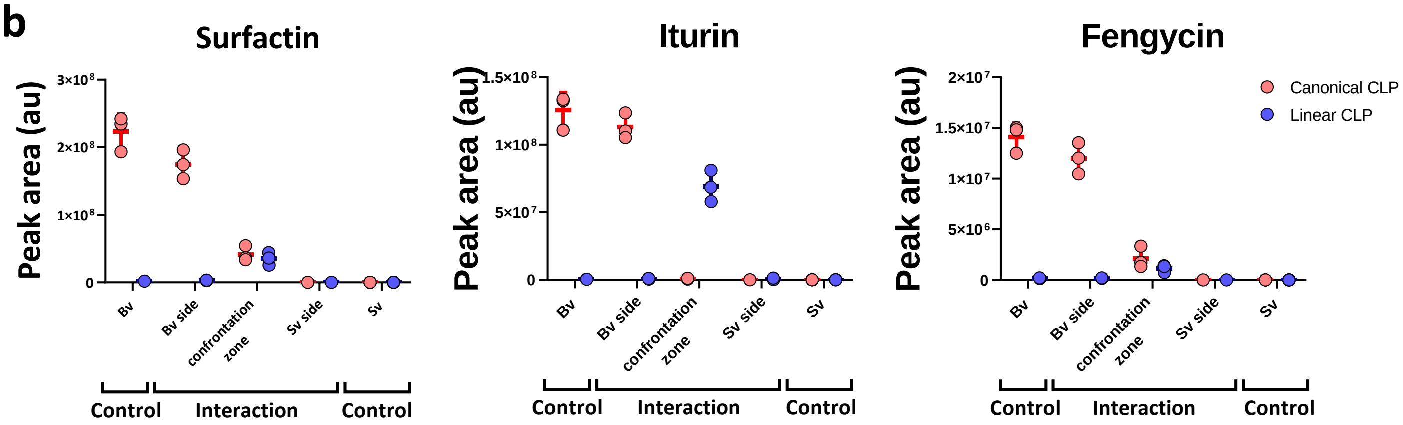

b
Surfactin
Iturin
Fengycin
Peak area (au)
3×10⁸
2×10⁸
1×10⁸
0
1.5×10⁸
1×10⁸
5×10⁷
2×10⁷
1.5×10⁷
1×10⁷
5×10⁶
Canonical CLP
Linear CLP
Bv
Bv side
confrontation zone
Sv side
Sv
Control
Interaction
Control

Supp. Fig. 2

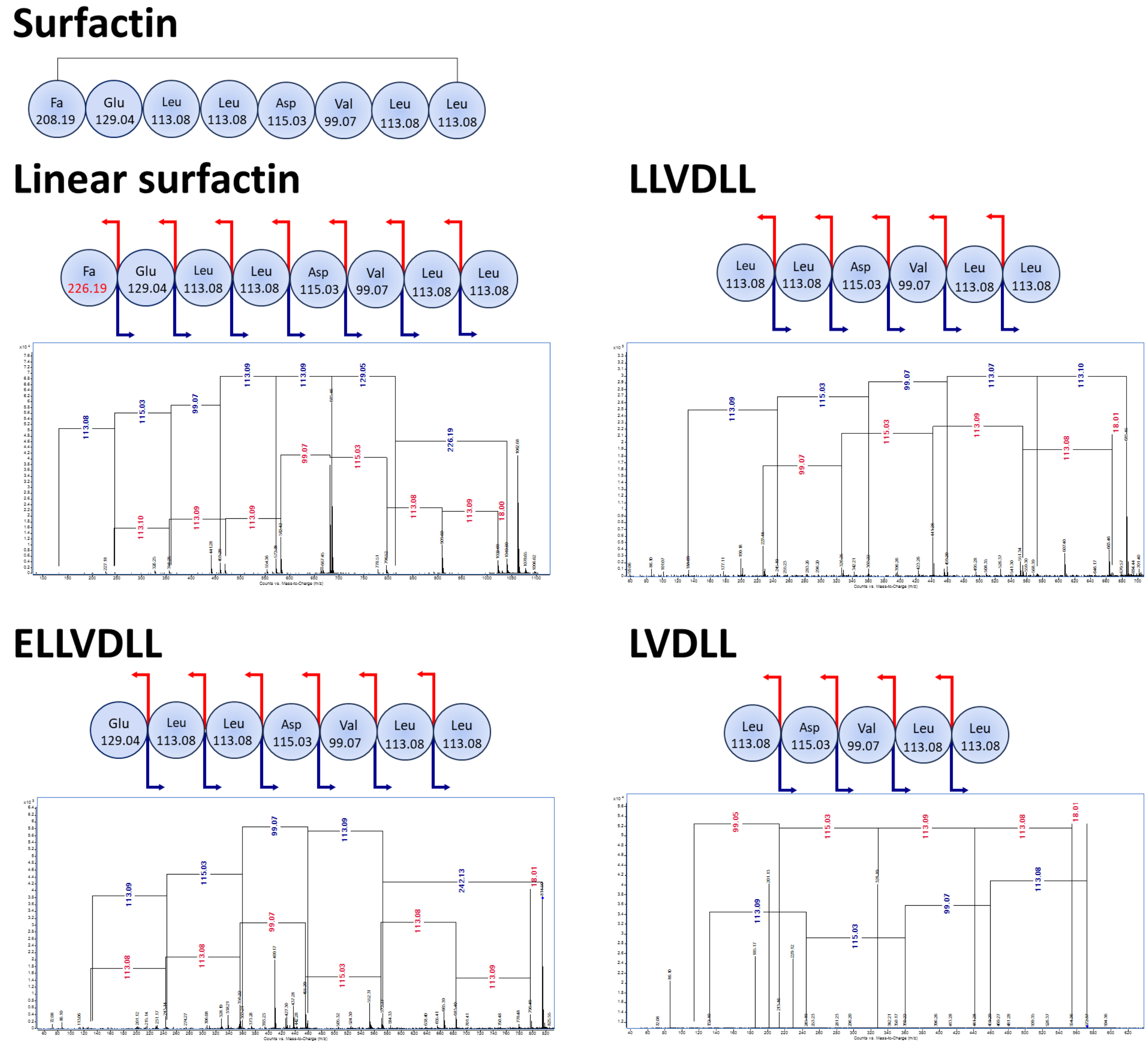
Surfactin
Fa 208.19
Glu 129.04
Leu 113.08
Leu 113.08
Asp 115.03
Val 99.07
Leu 113.08
Leu 113.08
Linear surfactin
Fa 226.19
Glu 129.04
Leu 113.08
Leu 113.08
Asp 115.03
Val 99.07
Leu 113.08
Leu 113.08
LLVDLL
Leu 113.08
Leu 113.08
Asp 115.03
Val 99.07
Leu 113.08
Leu 113.08
ELLVDLL
Glu 129.04
Leu 113.08
Leu 113.08
Asp 115.03
Val 99.07
Leu 113.08
Leu 113.08
LVDLL
Leu 113.08
Asp 115.03
Val 99.07
Leu 113.08
Leu 113.08

Supp. Fig. 3

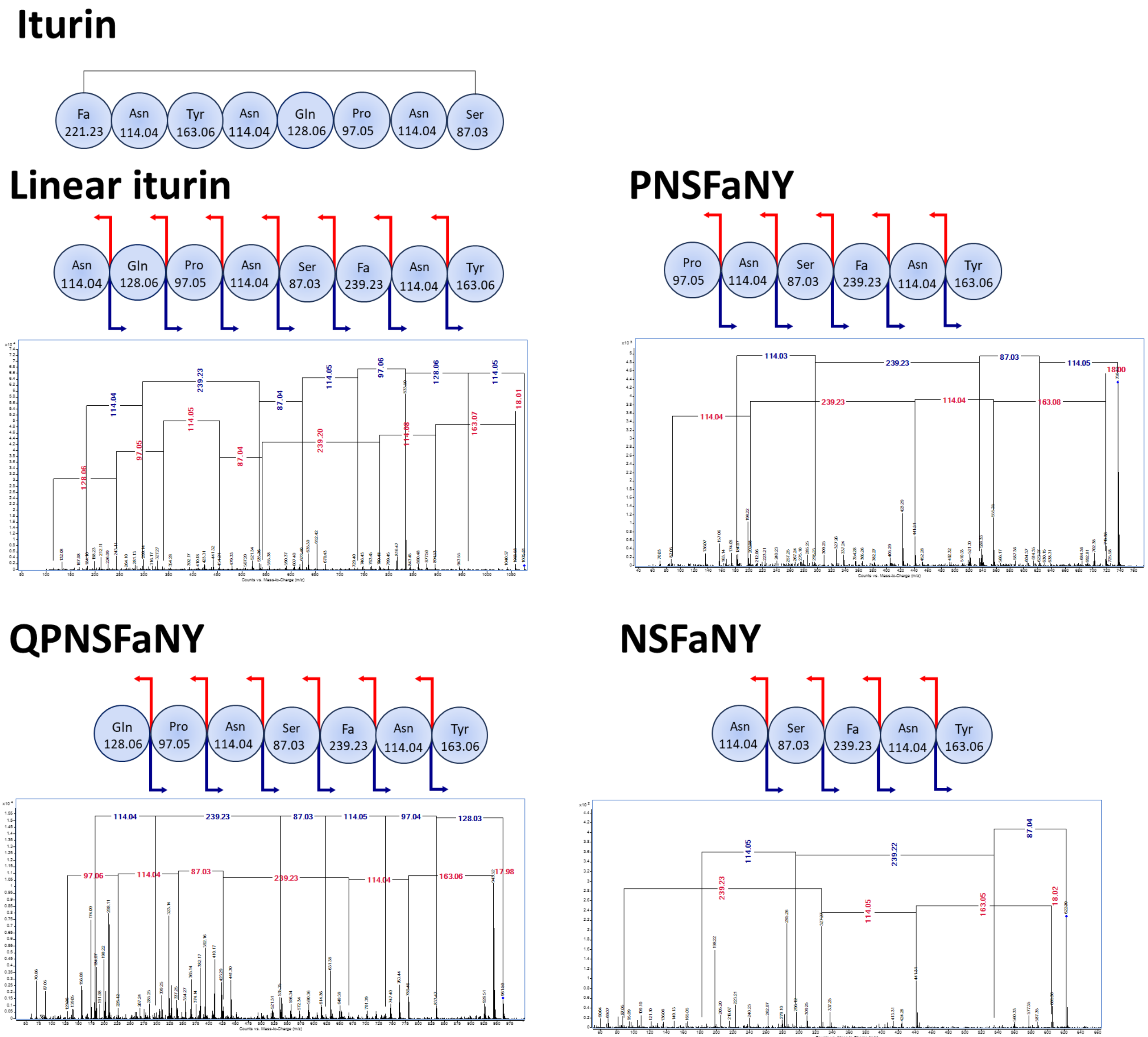

Iturin
Fa 221.23
Asn 114.04
Tyr 163.06
Asn 114.04
Gln 128.06
Pro 97.05
Asn 114.04
Ser 87.03
Linear iturin
Asn 114.04
Gln 128.06
Pro 97.05
Asn 114.04
Ser 87.03
Fa 239.23
Asn 114.04
Tyr 163.06
PNSFaNY
Pro 97.05
Asn 114.04
Ser 87.03
Fa 239.23
Asn 114.04
Tyr 163.06
QPNSFaNY
Gln 128.06
Pro 97.05
Asn 114.04
Ser 87.03
Fa 239.23
Asn 114.04
Tyr 163.06
NSFaNY
Asn 114.04
Ser 87.03
Fa 239.23
Asn 114.04
Tyr 163.06

**Supp. Fig. 4**

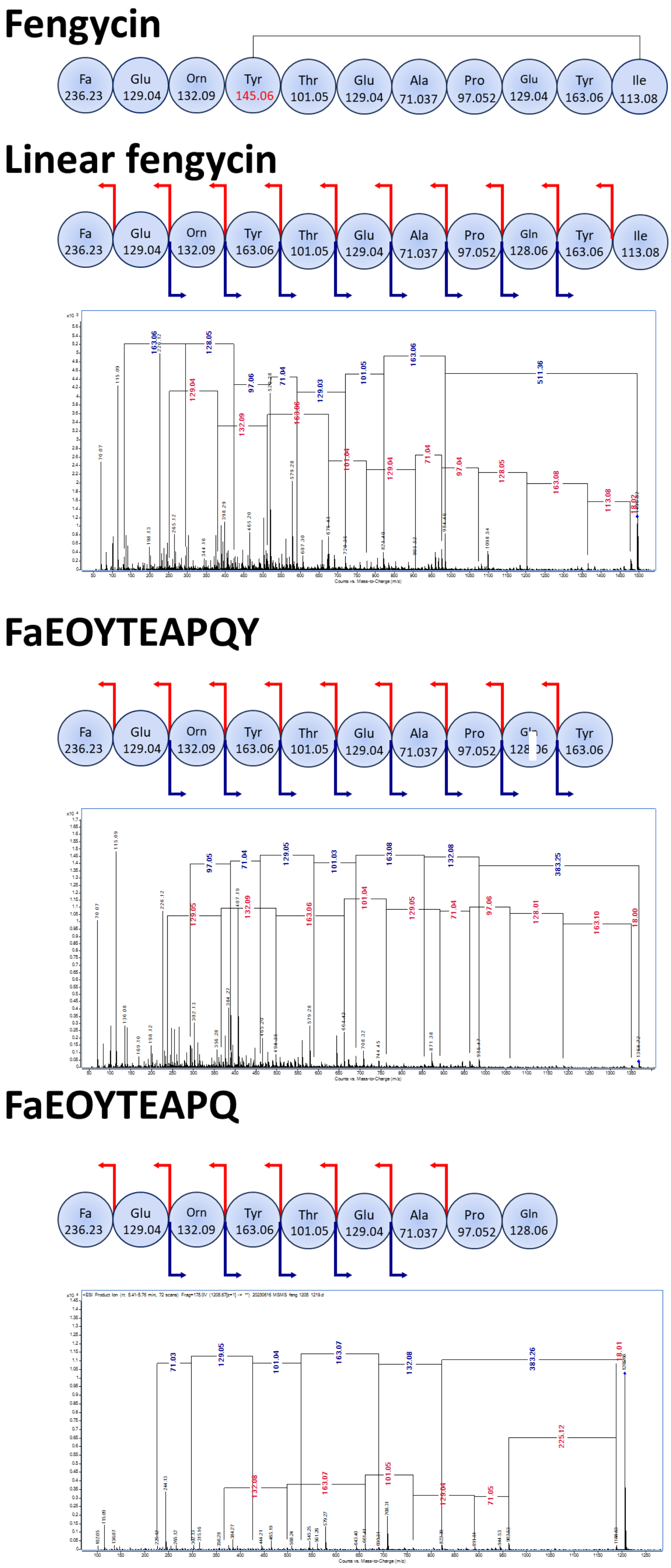

Fengycin
Fa 236.23
Glu 129.04
Orn 132.09
Tyr 145.06
Thr 101.05
Glu 129.04
Ala 71.037
Pro 97.052
Glu 129.04
Tyr 163.06
Ile 113.08
Linear fengycin
Fa 236.23
Glu 129.04
Orn 132.09
Tyr 163.06
Thr 101.05
Glu 129.04
Ala 71.037
Pro 97.052
Gln 128.06
Tyr 163.06
Ile 113.08
Counts vs. Mass-to-Charge (m/z)
FaEOYTEAPQY
Fa 236.23
Glu 129.04
Orn 132.09
Tyr 163.06
Thr 101.05
Glu 129.04
Ala 71.037
Pro 97.052
Tyr 163.06
Counts vs. Mass-to-Charge (m/z)
FaEOYTEAPQ
Fa 236.23
Glu 129.04
Orn 132.09
Tyr 163.06
Thr 101.05
Glu 129.04
Ala 71.037
Pro 97.052
Gln 128.06
Counts vs. Mass-to-Charge (m/z)

**Supp. Fig. 5**

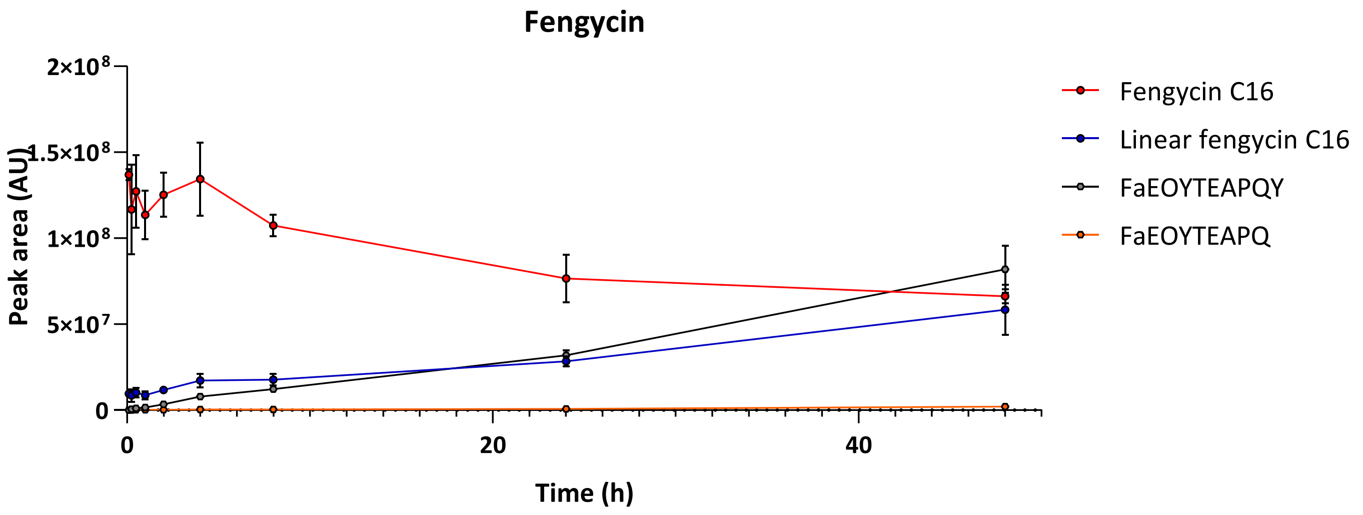

Fengycin
2×10⁸
1.5×10⁸
1×10⁸
5×10⁷
0
Peak area (AU)
0
20
40
Time (h)
Fengycin C16
Linear fengycin C16
FaEOYTEAPQY
FaEOYTEAPQ


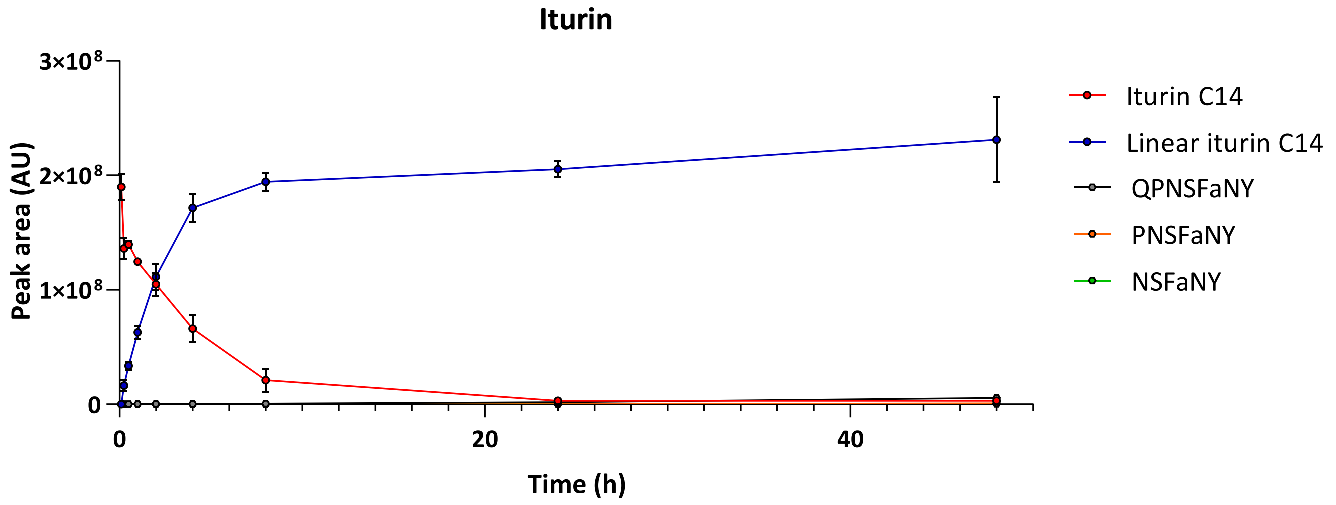

Iturin
3×10⁸
2×10⁸
1×10⁸
0
Peak area (AU)
0
20
40
Time (h)
Iturin C14
Linear iturin C14
QPNSFaNY
PNSFaNY
NSFaNY


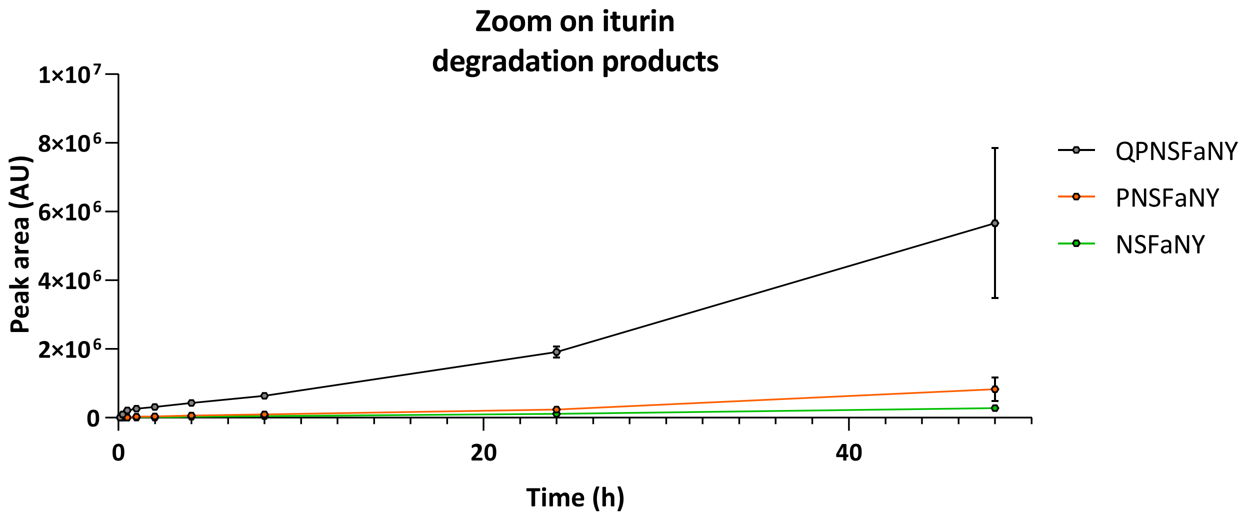

Zoom on iturin
degradation products
1×10⁷
8×10⁶
6×10⁶
4×10⁶
2×10⁶
0
Peak area (AU)
0
20
40
Time (h)
QPNSFaNY
PNSFaNY
NSFaNY

**Supp. Fig. 6**

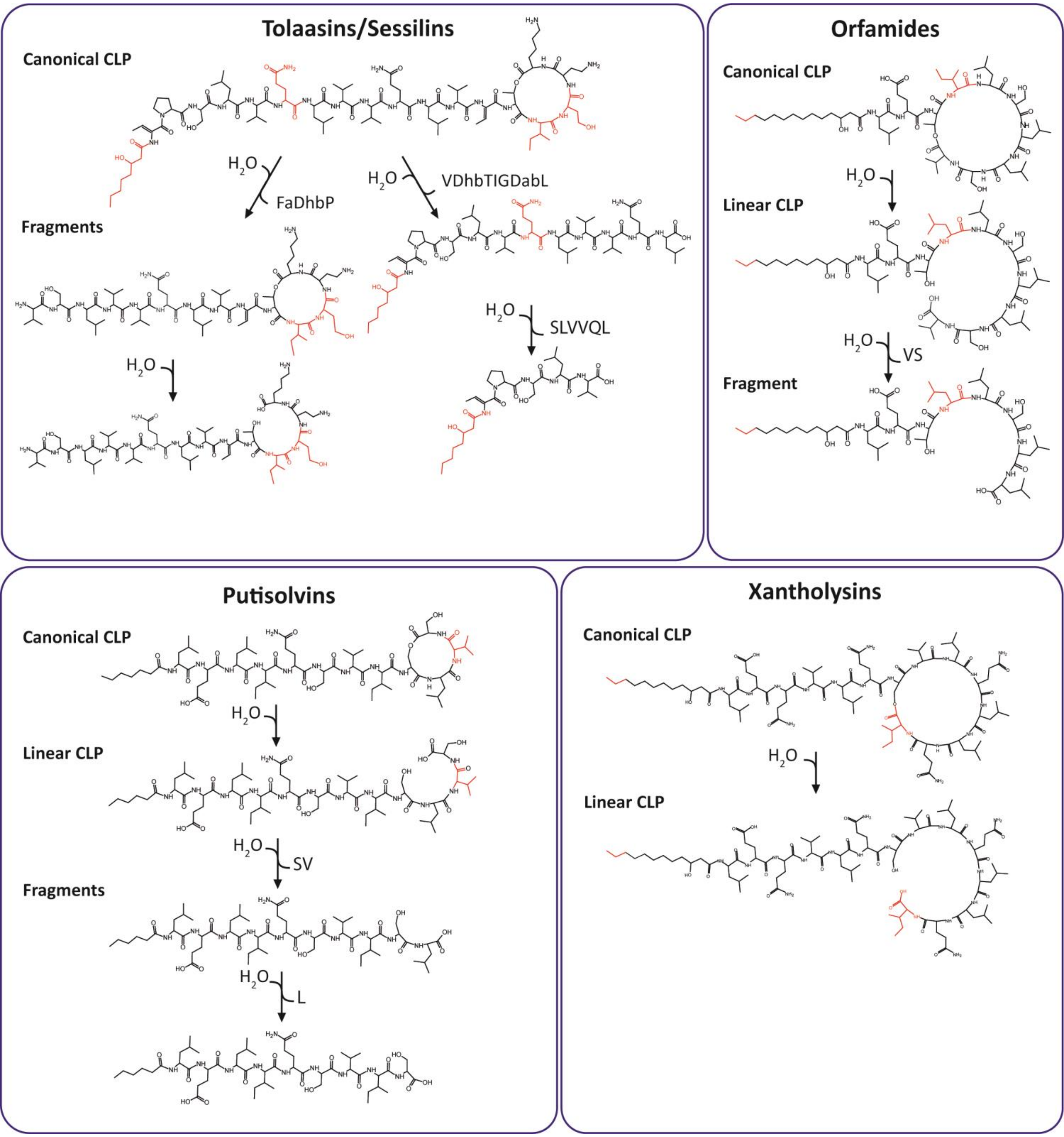

Tolaasins/Sessilins
Canonical CLP
$H_2O$
FaDhbP
$H_2O$
VDhbTIGDabL
Fragments
$H_2O$
$H_2O$
SLVVQL
Orfamides
Canonical CLP
$H_2O$
Linear CLP
$H_2O$
VS
Fragment
Putisolvins
Canonical CLP
$H_2O$
Linear CLP
$H_2O$
SV
Fragments
$H_2O$
L
Xantholysins
Canonical CLP
$H_2O$
Linear CLP

**Supp. Fig. 7**

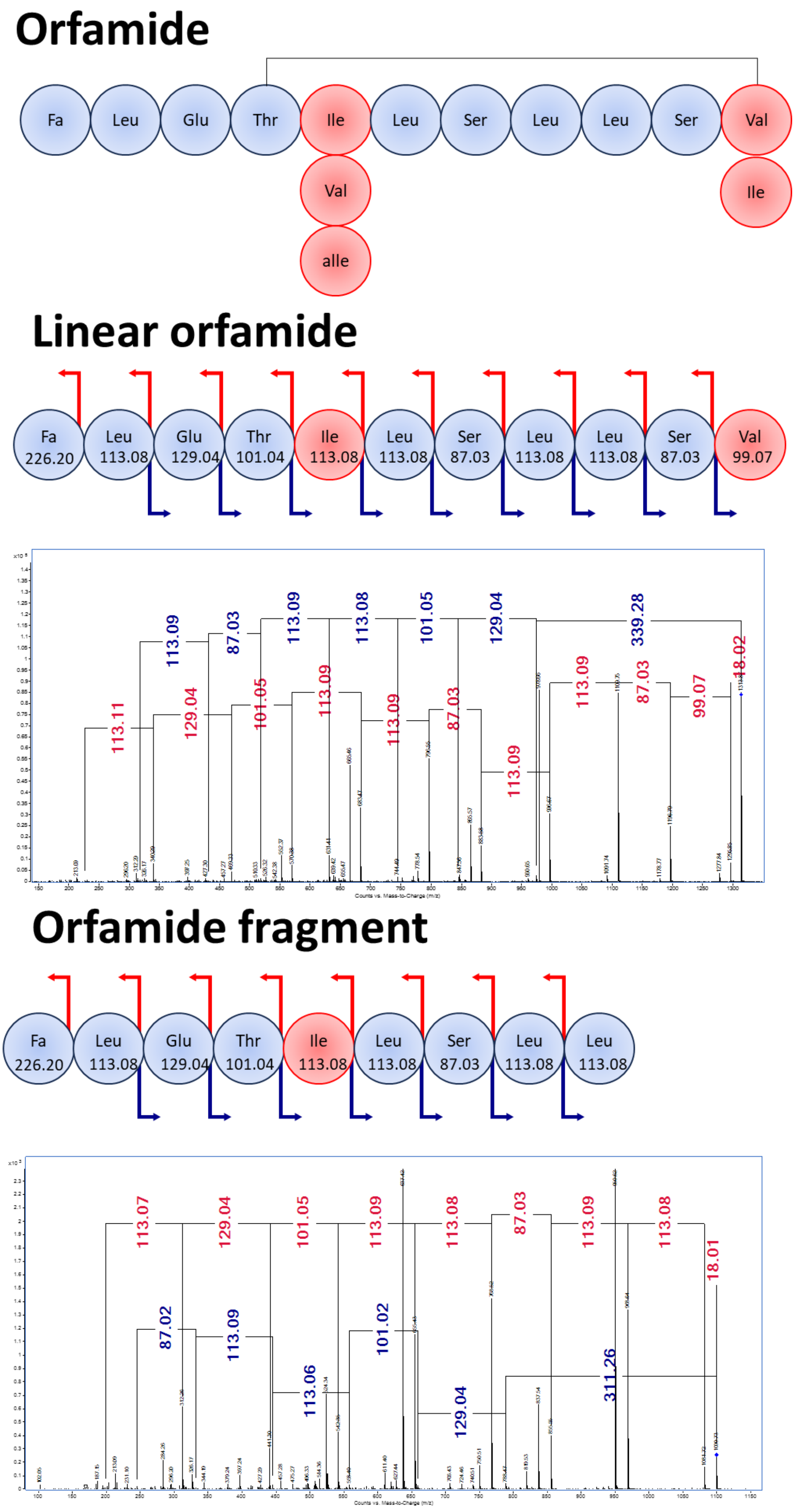
Orfamide
Fa
Leu
Glu
Thr
Ile
Leu
Ser
Leu
Leu
Ser
Val
Val
Ile
alle
Linear orfamide
Fa 226.20
Leu 113.08
Glu 129.04
Thr 101.04
Ile 113.08
Leu 113.08
Ser 87.03
Leu 113.08
Leu 113.08
Ser 87.03
Val 99.07
113.09
87.03
113.09
113.08
101.05
129.04
339.28
113.11
129.04
101.05
113.09
113.09
87.03
113.09
113.09
87.03
99.07
18.02
Counts vs. Mass-to-Charge (m/z)
Orfamide fragment
Fa 226.20
Leu 113.08
Glu 129.04
Thr 101.04
Ile 113.08
Leu 113.08
Ser 87.03
Leu 113.08
Leu 113.08
113.07
129.04
101.05
113.09
113.08
87.03
113.09
113.08
18.01
87.02
113.09
113.06
101.02
129.04
311.26
Counts vs. Mass-to-Charge (m/z)

Supp. Fig. 8

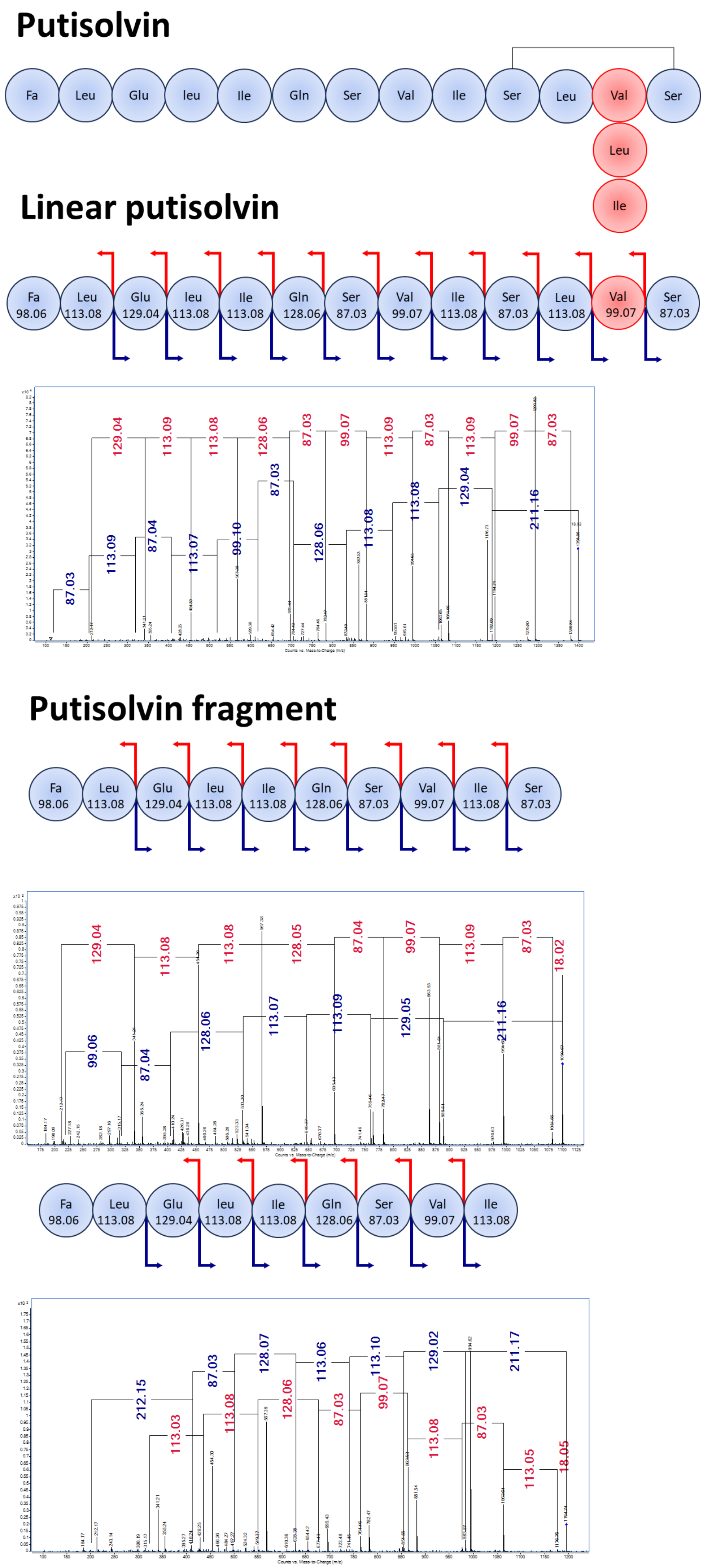

Putisolvin
Fa Leu Glu leu Ile Gln Ser Val Ile Ser Leu Val Ser
Leu
Ile
Linear putisolvin
Fa 98.06 Leu 113.08 Glu 129.04 leu 113.08 Ile 113.08 Gln 128.06 Ser 87.03 Val 99.07 Ile 113.08 Ser 87.03 Leu 113.08 Val 99.07 Ser 87.03
129.04 113.09 113.08 128.06 87.03 99.07 113.09 87.03 113.09 99.07 87.03
87.03 113.09 87.04 113.07 99.10 87.03 128.06 113.08 113.08 129.04 211.16
Putisolvin fragment
Fa 98.06 Leu 113.08 Glu 129.04 leu 113.08 Ile 113.08 Gln 128.06 Ser 87.03 Val 99.07 Ile 113.08 Ser 87.03
129.04 113.08 113.08 128.05 87.04 99.07 113.09 87.03 18.02
99.06 87.04 128.06 113.07 113.09 129.05 211.16
Fa 98.06 Leu 113.08 Glu 129.04 leu 113.08 Ile 113.08 Gln 128.06 Ser 87.03 Val 99.07 Ile 113.08
212.15 87.03 128.07 113.06 113.10 129.02 211.17
113.03 113.08 128.06 87.03 99.07 113.08 87.03 113.05 18.05
Counts vs. Mass-to-Charge (m/z)

**Supp. Fig. 9**

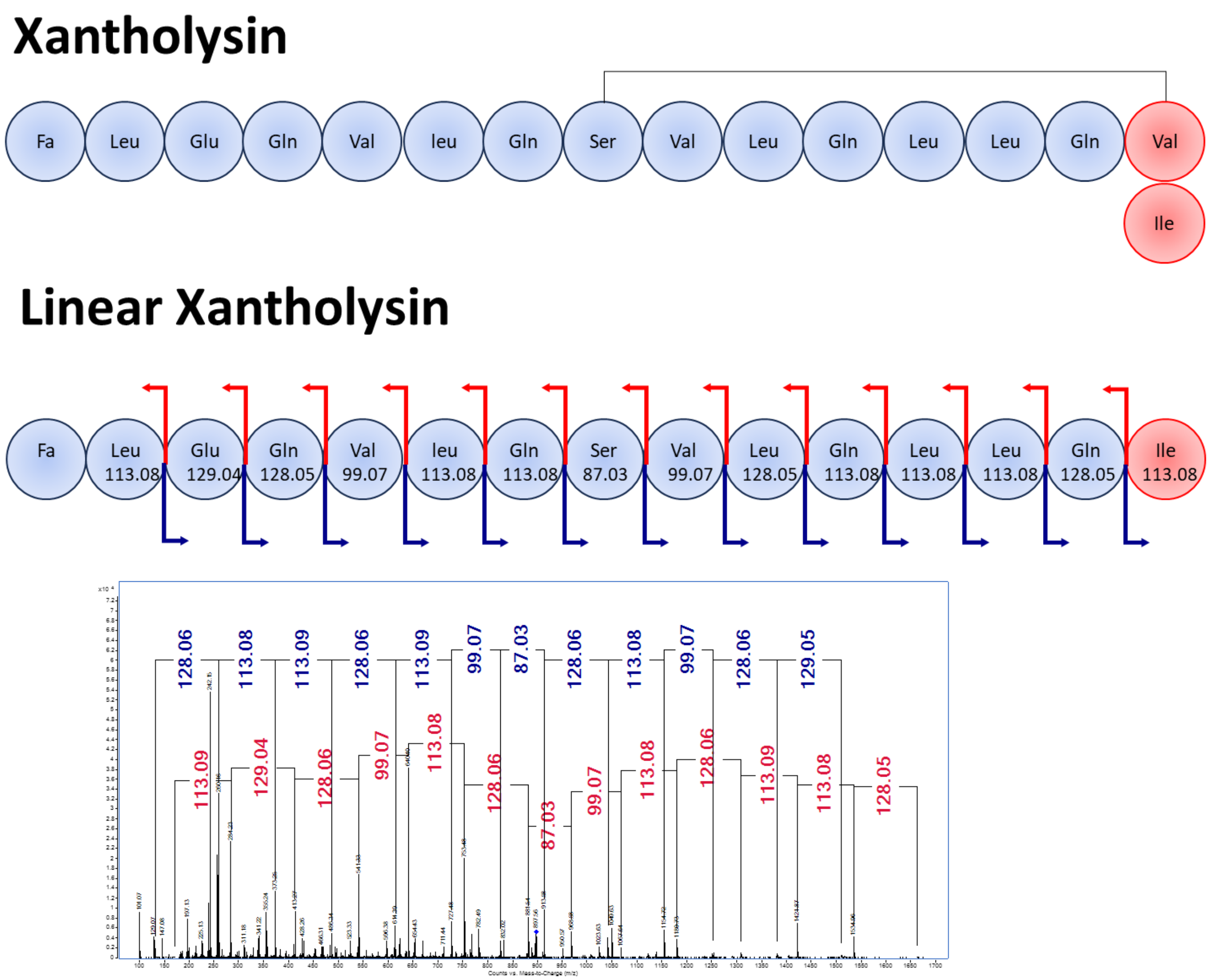

Xantholysin
Fa
Leu
Glu
Gln
Val
leu
Gln
Ser
Val
Leu
Gln
Leu
Leu
Gln
Val
Ile
Linear Xantholysin
Fa
Leu 113.08
Glu 129.04
Gln 128.05
Val 99.07
leu 113.08
Gln 113.08
Ser 87.03
Val 99.07
Leu 128.05
Gln 113.08
Leu 113.08
Leu 113.08
Gln 128.05
Ile 113.08
128.06
113.08
113.09
128.06
113.09
99.07
87.03
128.06
113.08
99.07
128.06
129.05
113.09
129.04
128.06
99.07
113.08
128.06
87.03
99.07
113.08
128.06
113.09
113.08
128.05

### Supp. Fig. 10

#### Sessilin/Tolaasin

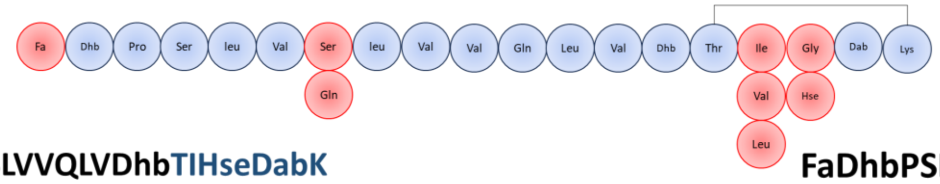


#### VSLVVQLVDhbTIHseDabK

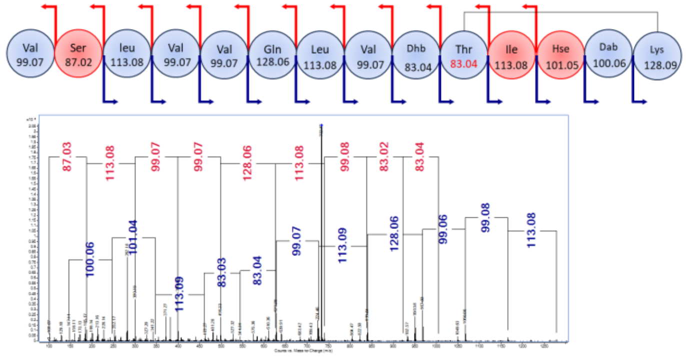


#### FaDhbPSLVQLVVQL

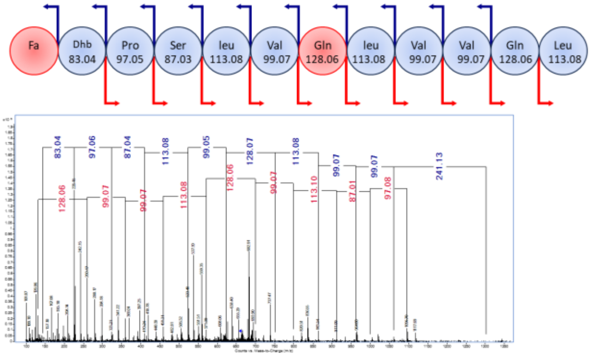


#### VSLVVQLVDhbTIHseDabK

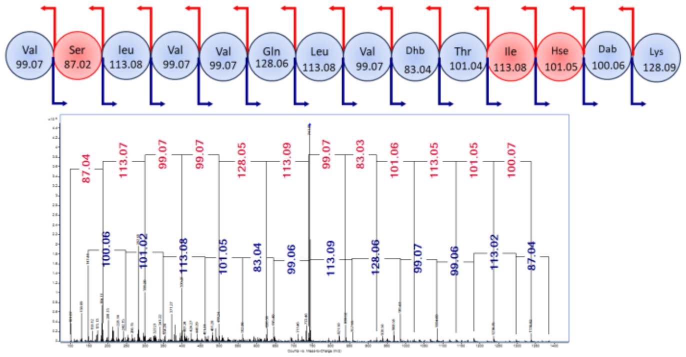


#### FaDhbPSLV

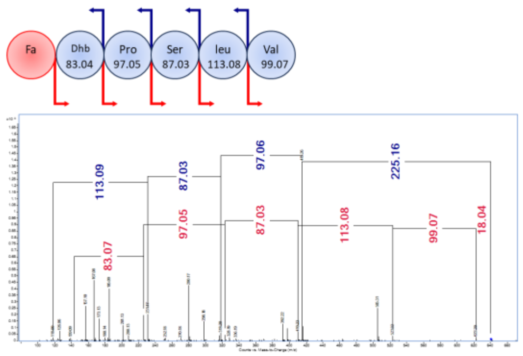

### Supp. Fig. 11

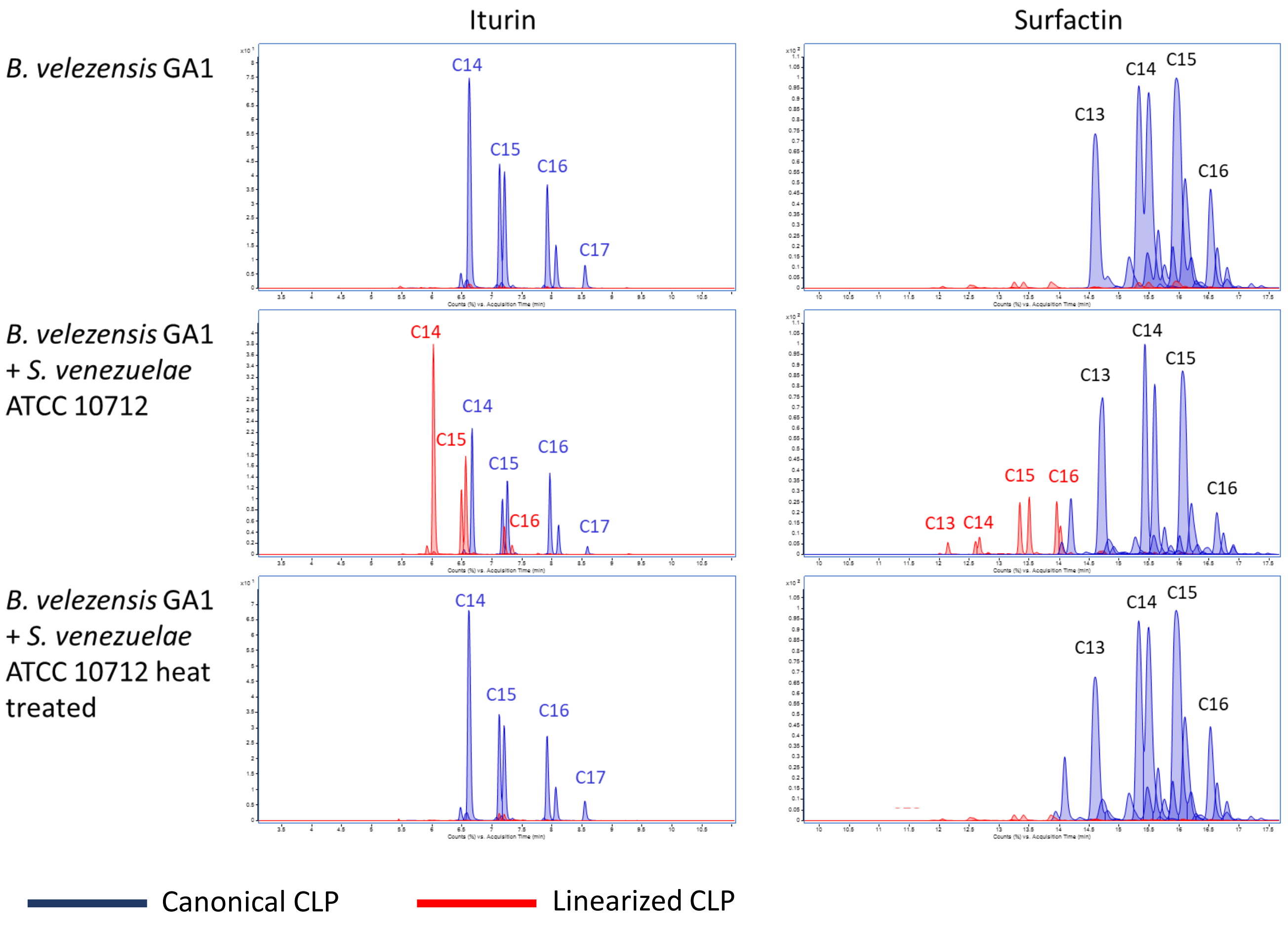
Iturin
Surfactin
*B. velezensis* GA1
*B. velezensis* GA1 + *S. venezuelae* ATCC 10712
*B. velezensis* GA1 + *S. venezuelae* ATCC 10712 heat treated
C13
C14
C15
C16
C17
Canonical CLP
Linearized CLP

**Supp. Fig. 12**

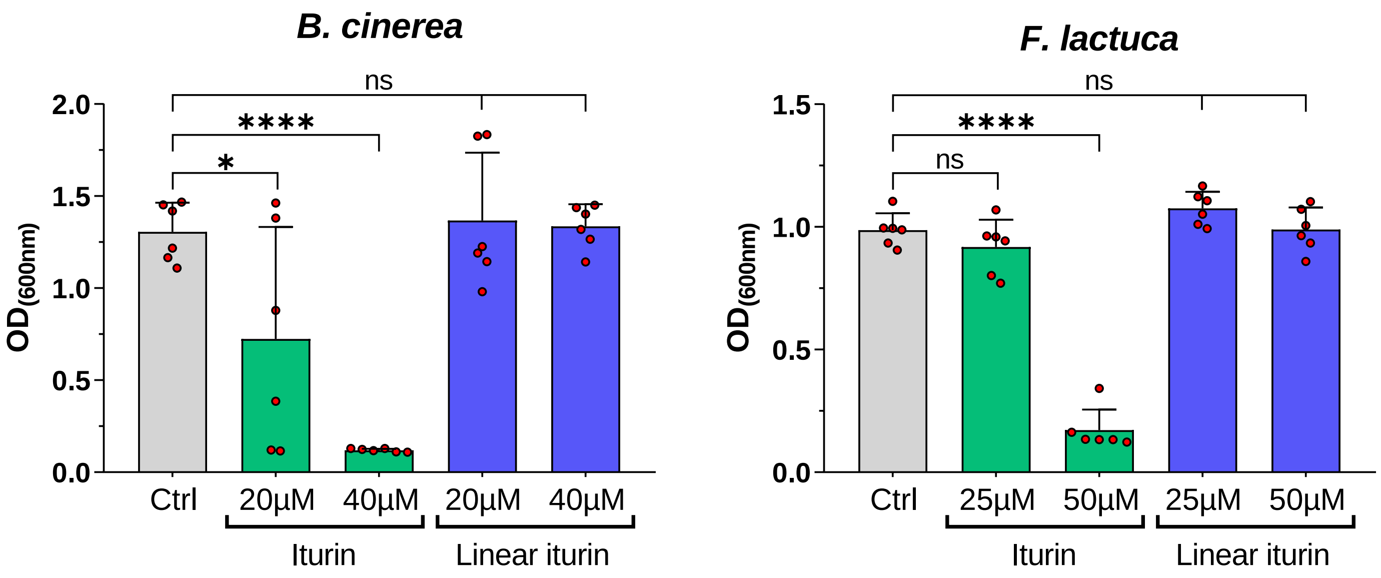

B. cinerea
OD(600nm)
2.0
1.5
1.0
0.5
0.0
ns
****
*
Ctrl
20µM
40µM
20µM
40µM
Iturin
Linear iturin
F. lactuca
OD(600nm)
1.5
1.0
0.5
0.0
ns
****
ns
Ctrl
25µM
50µM
25µM
50µM
Iturin
Linear iturin

#### Supp. Table 1

| ***Bacillus* strains** | **Locus-tag(s) of mutation** | **Characteristics** | **Relevant phenotype** | **Sources** |
|---|---|---|---|---|
| *B. velezensis* GA1 | | Wild-type strain | | Arguelles-Arias *et al.* 2008 |
| *B. velezensis* GA1 Δ*sfp* | GL331_06785 | Δ*sfp* ::*chl* ; Chl+ | GA1 deleted of the 4'phosphopantetheinyl transferase *sfp* gene; unable to produce lipopeptides, polyketides and bacillibactin | Andric *et al.,* 2022 |
| *B. velezensis* GA1 Δ*baeJ* Δ*dfnA* Δ*mlnA* | GL331_13510-15925-12195 | Δ*baeJ* ::*chl* Δ*dfnA::phl* Δ*mlnA* ::neo ; Chl+ Phl+ Neo+ | GA1 deleted of *baeJ*, *dfnM* and *mlnA* genes; unable to produce bacillaenes, difficidins and macrolactins | This study |

| ***Streptomyces* strains** | | **Characteristics** | **Relevant phenotype** | **Sources** |
|---|---|---|---|---|
| *S. venezuelae* ATCC 10712 | | Wild-type strain | | Rigali Laboratory |

| ***Pseudomonas* strains** | | | | |
|---|---|---|---|---|
| *P. mosselii* BW11M1 | | Wild-type strain | Xantholysins producer | Andric *et al.*, 2021 |
| *P. tolaasii* CH36 | | Wild-type strain | Tolaasins producer | Andric *et al.*, 2021 |
| *P. sessilinigenes* CMR12a | | Wild-type strain | Sessilins and Orfamides producer | Andric *et al.*, 2021 |
| *P. protegens* Pf-5 | | Wild-type strain | Orfamides producer | Andric *et al.*, 2021 |
| *P. lactis* SS101 | | Wild-type strain | Massetolides producer | Andric *et al.*, 2021 |
| *P. putida* WCU64 | | Wild-type strain | Putisolvins producer | Andric *et al.*, 2021 |

| **Phytopathogenic strains** | | | | |
|---|---|---|---|---|
| *Fusarium lactuca* F 1.39 | | Wild-type strain | Fusariose agent | Höfte Laboratory |
| *Botytis cinerea* R16 | | Wild-type strain | grey mould disease agent | Höfte Laboratory |

### Supp. Table 2

| Step | Parameter | | | | |
|---|---|---|---|---|---|
| **Step 0** | **Parameter** | | | | |
| **Data export to mz-mine** | **export** | as mzdata | entire data file | | |
| | **MS level** | All | | | |
| | **MS storage** | centroid data | | | |
| | **compute deisotope** | ND | | | |
| | **Max spike width** | 2 | | | |
| | **Required valley** | 0.7 | | | |
| | **Height filters** | Absolute height | >=2500 counts | | |
| | **Max number of peaks** | limit to the largest | ND | | |
| | **Charge state** | ND | | | |
| MzMine Prameters | | | | | |
| **Step 1** | **Parameter** | | | | |
| **Mass detection** | | | | | |
| | **MS level** | 1 | | | |
| | | noize level | 1500 | | |
| | | Detect isotope below noise level | TRUE | | |
| | | Chemical elements | H,C,N,O,S | | |
| | m/z tolerence | m/z | 0.002 | ppm | 5 |
| | **Ms level** | 2 | | | |
| | | noize level | nd | | |
| | | Detect isotope below noise level | FALSE | | |
| **Step 2** | **Parameter** | | | | |
| **ADAP chromatogram builder** | | | | | |
| | **Scans** | MS level (1) | | | |
| | **Min group size in #scans** | 3 | | | |
| | **Group intensity threshold** | 5000 | | | |
| | **Min highest intensity** | 7500 | | | |
| | **m/z tolerence** | m/z | 0.01 | ppm | 25 |
| **Step 3** | **Parameter** | | | | |
| **Chromatogram deconvalution** | | | | | |
| | **Algorithm** | **Local minimum search** | | | |
| | | Dimension | Rentetion time | | |
| | | Chromatographic threshold | 85% | | |
| | | Serach min Rt range (min) | 0.2 | | |
| | | Min relative height | 0.1 | | |
| | | Min absolute height | 6000 | | |
| | | Min ratio of peak top/edge | 1.50 | | |
| | | Peak duration range (min) | 0-1.0 | | |
| | | Min # of data points | 3 | | |
| | **m/z center calculation** | Median | | | |
| | **m/z range for MS2 scan pairing (DA)** | TRUE | | | |
| | **Rt range for MS2 scan pairing (min)** | 0.2 | | | |
| | **MS1 to MS2 precursor tolerance (m/z)** | m/z | 0.02 | ppm | 20 |
| **Step 4** | **Parameter** | | | | |
| **Isotope peak grouper** | **mz tolerance** | mz | 0 | ppm | 15 |
| | **Retention time** | 0.15 | | | |
| | **Monotonic shape** | nd | | | |
| | **Maximum charge** | 2 | | | |
| | **Representative isotope** | lowest m/z | | | |
| **Step 5** | **Parameter** | | | | |
| **Join aligner** | | | | | |
| | **m/z tolerance** | m/z | 0 | ppm | 25 |
| | **weight for m/z** | 75 | | | |
| | **Retention time tolerance** | 0.2 min | | | |
| | **Weight for RT** | 25 | | | |
| | **Require same chare state** | FALSE | | | |
| | **Compare isotope pattern** | FALSE | | | |
| | **Compare spectra similarity** | FALSE | | | |
| **Step 6** | **Parameter** | | | | |
| **Duplicate list filter** | | | | | |
| | **m/z tolerance** | m/z | 0 | ppm | 25 |
| | **filter mode** | New average | | | |
| | **Rt tolerance** | 0.02 absolute (min) | | | |
| | **Require same identification** | FALSE | | | |
| **Step 6** | **Parameter** | | | | |
| **Feature list filter** | | | | | |
| | **minimum peaks in a row** | FALSE | | | |
| | **minimum peaks in an isotope pattern** | 2 | | | |
| | **Validate 13C isotope pattern** | FALSE | | | |
| | **m/z** | 350.01-1125.0 | | | |
| | **Rt (min)** | 0.5-25.0 | | | |
| | **Peak duration range** | ND | | | |
| | **Chromatographic FWHM** | ND | | | |
| | **Charge** | FALSE | | | |
| | **Kendrick mass defect** | FALSE | | | |
| | **Parameter** | No parameter defined | | | |
| | **Only identified?** | FALSE | | | |
| | **Test in identity** | FALSE | | | |
| | **Text in comment** | FALSE | | | |
| | **Keep or remove rows** | Keep rows that match all criteria | | | |
| | **Keep only peaks with MS2 scan (GNPS)** | FALSE | | | |
| | **Reset the peak number ID** | FALSE | | | |
| **Step 6** | **Parameter** | | | | |
| **Gap filling** | **Intensity tolerance** | 0.05 | | | |
| | **m/z tolerance** | 0.025 | m/z | 35 | ppm |
| | **Rt tolerance** | 0.1 | | | |
| | **Minimum data points** | 3 | | | |

### Supp. Table. 3

<table>
<tr><th>T: Protein IDs</th><th>T: Majority protein IDs</th><th>Function</th><th>Function type</th></tr>
<tr><td>tr|F2R101|F2R101_STRVP</td><td>tr|F2R101|F2R101_STRVP</td><td>Putative secreted peptidase</td><td>Protease/peptidase</td></tr>
<tr><td>tr|F2RJA5|F2RJA5_STRVP</td><td>tr|F2RJA5|F2RJA5_STRVP</td><td>Secreted protease</td><td></td></tr>
<tr><td>tr|F2R8T0|F2R8T0_STRVP</td><td>tr|F2R8T0|F2R8T0_STRVP</td><td>Putative hydrolase</td><td></td></tr>
<tr><td>tr|F2RB07|F2RB07_STRVP</td><td>tr|F2RB07|F2RB07_STRVP</td><td>Peptidase S8 and S53, subtilisin, kexin,sedolisin</td><td></td></tr>
<tr><td>tr|F2RBP4|F2RBP4_STRVP</td><td>tr|F2RBP4|F2RBP4_STRVP</td><td>Peptidase M23 domain-containing protein</td><td></td></tr>
<tr><td>tr|F2RES2|F2RES2_STRVP</td><td>tr|F2RES2|F2RES2_STRVP</td><td>Putative peptidase</td><td></td></tr>
<tr><td>tr|F2RGW0|F2RGW0_STRVP</td><td>tr|F2RGW0|F2RGW0_STRVP</td><td>Branched-chain amino acid ABC transporter, amino acid-binding protein</td><td>Amino acid/peptide transport</td></tr>
<tr><td>tr|F2R1F6|F2R1F6_STRVP</td><td>tr|F2R1F6|F2R1F6_STRVP</td><td>Dipeptide transport system permease protein DppC</td><td></td></tr>
<tr><td>tr|F2R1F7|F2R1F7_STRVP;tr|F2R1G3|F2R1G3_STRVP</td><td>tr|F2R1F7|F2R1F7_STRVP</td><td>Oligopeptide transport ATP-binding protein OppD</td><td></td></tr>
<tr><td>tr|F2R1F8|F2R1F8_STRVP;tr|F2R1G4|F2R1G4_STRVP</td><td>tr|F2R1F8|F2R1F8_STRVP</td><td>Oligopeptide transport ATP-binding protein OppF</td><td></td></tr>
<tr><td>tr|F2RCI4|F2RCI4_STRVP</td><td>tr|F2RCI4|F2RCI4_STRVP</td><td>Putative amino acid transporter</td><td></td></tr>
<tr><td>tr|F2R288|F2R288_STRVP</td><td>tr|F2R288|F2R288_STRVP</td><td>Fatty acid-binding protein DegV</td><td>Fatty acid transport</td></tr>
</table>

### Supp. Table. 4

**Surfactin features identified**

| m/z | Compound | Adduct |
|---|---|---|
| 994.65 | Surfactin C12 | $[m+H]^+$ |
| 1008.66 | Surfactin C13 | $[m+H]^+$ |
| 1022.68 | Surfactin C14 | $[m+H]^+$ |
| 1036.69 | Surfactin C15 | $[m+H]^+$ |
| 1050.71 | Surfactin C16 | $[m+H]^+$ |
| 1012.65 | Linear surfactin C12 | $[m+H]^+$ |
| 1026.67 | Linear surfactin C13 | $[m+H]^+$ |
| 1040.68 | Linear surfactin C14 | $[m+H]^+$ |
| 1054.7 | Linear surfactin C15 | $[m+H]^+$ |
| 1068.72 | Linear surfactin C16 | $[m+H]^+$ |
| 814.49 | LLVALL | $[m+H]^+$ |
| 685.45 | LLVAL | $[m+H]^+$ |
| 699.47 | LLLAL | $[m+H]^+$ |
| 572.37 | LLVA | $[m+H]^+$ |
| 586.38 | LLLA | $[m+H]^+$ |
| 360.24 | C13-E | $[m+H]^+$ |
| 374.25 | C14-E | $[m+H]^+$ |
| 388.25 | C15-E | $[m+H]^+$ |
| 402.28 | C16-E | $[m+H]^+$ |
| 487.34 | C15-EL | $[m+H]^+$ |
| 501.35 | C16-EL | $[m+H]^+$ |
| 711.47 | Not identified | [m+H]+ |
| 598.38 | Not identified | [m+H]+ |

**Iturin features identified**

| m/z | Compound | Adduct |
|---|---|---|
| 1001.53 | Iturin A C11 | $[m+H]^+$ |
| 1015.53 | Iturin A C12 | $[m+H]^+$ |
| 1029.53 | Iturin A C13 | $[m+H]^+$ |
| 1043.54 | Iturin A C14 | $[m+H]^+$ |
| 1044.54 | Iturin C C14 | $[m+H]^+$ |
| 1057.56 | Iturin A C15 | $[m+H]^+$ |
| 1071.57 | Iturin A C16 | $[m+H]^+$ |
| 1085.60 | Iturin A C17 | $[m+H]^+$ |
| 1019.52 | Linear iturin A C11 | $[m+H]^+$ |
| 1033.53 | Linear iturin A C12 | $[m+H]^+$ |
| 1047.55 | Linear iturin A C13 | $[m+H]^+$ |
| 1061.57 | Linear iturin A C14 | $[m+H]^+$ |
| 1075.57 | Linear iturin A C15 | $[m+H]^+$ |
| 1089.59 | Linear iturin A C16 | $[m+H]^+$ |
| 1103.61 | Linear iturin A C17 | $[m+H]^+$ |
| 947.51 | QPNS-C14-NY | $[m+H]^+$ |
| 961.53 | QPNS-C15-NY | $[m+H]^+$ |
| 975.55 | QPNS-C16-NY | $[m+H]^+$ |
| 819.46 | PNS-C14-NY | $[m+H]^+$ |
| 833.47 | PNS-C15-NY | $[m+H]^+$ |
| 736.42 | NS-C15-NY | $[m+H]^+$ |
| 1115.60 | Not identified | [m+H]+ |
| 1045.57 | Not identified | $[m+H]^+$ |
| 1073.56 | Not identified | $[m+H]^+$ |
| 1087.58 | Not identified | $[m+H]^+$ |
| 1091.57 | Not identified | $[m+H]^+$ |
| 930.49 | Not identified | $[m+H]^+$ |

**Fengycin features identified**

| m/z | Compound | Adduct |
|---|---|---|
| 1435.76 | Fengycin A C14 | $[m+H]^+$ |
| 1463.80 | Fengycin A C16 | $[m+H]^+$ |
| 1477.82 | Fengycin A C17 | $[m+H]^+$ |
| 1491.83 | Fengycin A C18 | $[m+H]^+$ |
| 1505.85 | Fengycin A C19 | $[m+H]^+$ |
| 1453.78 | Linear fengycin A C14 | $[m+H]^+$ |
| 1467.79 | Linear fengycin A C15 | $[m+H]^+$ |
| 1481.81 | Linear fengycin A C16 | $[m+H]^+$ |
| 1482.8 | Linear fengycin A C16-$Glu^8$ | $[m+H]^+$ |
| 1495.83 | Linear fengycin A C17 | $[m+H]^+$ |
| 1509.84 | Linear fengycin A C18 | $[m+H]^+$ |
| 1510.83 | Linear fengycin A C18-$Glu^8$ | $[m+H]^+$ |
| 1523.85 | Linear fengycin A C19 | $[m+H]^+$ |
| 1354.71 | C15-EOYTEAPQY | $[m+H]^+$ |
| 1368.73 | C16-EOYTEAPQY | $[m+H]^+$ |
| 1382.74 | C17-EOYTEAPQY | $[m+H]^+$ |
| 1396.76 | C18-EOYTEAPQY | $[m+H]^+$ |
| 1410.77 | C19-EOYTEAPQY | $[m+H]^+$ |
| 1424.79 | C20-EOYTEAPQY | $[m+H]^+$ |
| 1219.68 | C17-EOYTEAPQY | $[m+H]^+$ |
